# State-dependent release of extracellular particles with distinct α2,6-sialylation patterns and small RNA cargo related to neuroinflammation

**DOI:** 10.1101/2025.10.30.684535

**Authors:** Marta Garcia-Contreras, Carinna Lima, Eric Alsop, Balazs Kaszala, Bessie Meechoovet, Benjamin Purnell, Nan Jiang, Andras Saftics, Candice Tang, Olivia D Walsh, Sophia Holland, Jimmy Fay, Barbara Smith, Krishna P Sigdel, Kendall Van Keuren-Jensen, Tijana Jovanovic-Talisman, Saumya Das

**Affiliations:** Cardiovascular Research Center, Massachusetts General Hospital, Harvard Medical School, Boston, MA, USA; Department of Cancer Biology and Molecular Medicine, Beckman Research Institute, City of Hope Comprehensive Cancer Center, Duarte, CA, USA; Translational Genomics Research Institute, Phoenix, AZ, USA; Department of Physics and Astronomy, California State Polytechnic University, Pomona, CA, USA; NanoFCM, ltd, Nottingham, UK; Department of Cell Biology, Johns Hopkins University, Baltimore, MD, USA

**Keywords:** Extracellular vesicles, exomeres, supermeres, microglia, neuroinflammation, miRNA

## Abstract

Neuroinflammation is a significant contributor to neurodegenerative diseases, including Alzheimer’s disease, Parkinson’s disease, and related dementias; yet peripheral biomarkers for neuroinflammation remain an unmet medical need. Microglia, the resident immune cells of the central nervous system, play a dual role in maintaining homeostasis under physiological conditions and driving neuronal damage when chronically dysregulated. One mechanism by which microglia influence their environment is through the release of extracellular vesicles (EVs) and non-vesicular extracellular particles (NVEPs), which can serve as biomarkers the reflect cellular states. Here, we systematically isolated and characterized microglia-derived EVs and NVEPs under pro- and anti-inflammatory conditions and profiled their small RNA cargo by small RNA sequencing. We validated these findings in human iPSC-derived microglia and further recapitulated them in EVs and NVEPs from mouse brain and plasma. Using an engineered mouse model, we were able to isolate plasma microglia-specific EVs *in vivo* and demonstrated that their RNA cargo reflects their inflammatory state. Importantly, microglial EVs and NVEPs display distinct α2,6-sialylation patterns and small RNA signatures implicated in neurological diseases. These findings demonstrate that microglia-derived EVs and NVEPs cargo reflect microglial cellular state and establish them as putative minimally non-invasive biomarkers of early-stage neurodegenerative diseases.

## Introduction

Neuroinflammation is strongly associated with the onset and progression of various neurodegenerative diseases, including Alzheimer’s disease (AD), Parkinson’s disease (PD) and various forms of dementia^1,2^. Inflammatory acute injuries or disorders of the central nervous system (CNS) are either characterized by inflammation that leads to neurodegeneration (spinal cord injury, brain trauma, and stroke), or by neurodegeneration that induces inflammation (AD, PD, epilepsy, and Huntington’s disease). Among the key players in this process, microglia, the resident macrophage in the brain, has emerged as a key contributor. Under physiological conditions, microglia play a critical role in maintaining CNS homeostasis by synaptic pruning, efferocytosis, and phagocytosis of apoptotic cells and debris^3^. However, when chronically dysregulated, microglia can shift towards a pro-inflammatory phenotype characterized by secretion of both proteins, inflammatory cytokines, and chemokines as well as extracellular particles (EPs) into its local microenvironment, contributing to neuronal damage and accelerating disease progression^4–9^. Notably, neuroinflammation appears to be an early step in the pathogenesis of neurodegenerative diseases^10^. Accordingly, biomarkers to detect neuroinflammation would allow for identification of at-risk patients and early initiation of therapy.

EPs are secreted from most cell types and have emerged as critical mediators of intercellular communication in both essential physiology and pathogenesis of different diseases^11–14^. EPs are a heterogenous group of particles with a wide range of sizes (20-10,000 nm), cargo, and function; they have considerable potential as biomarkers for a range of diseases. The development of new isolation methods to improve the characterization of these varieties of EPs have led to the identification of new classes of extracellular particles, which are now classified as extracellular vesicles (EVs) and non-vesicular extracellular particles (NVEPs)^15^, although their detailed characterization in relevant human biofluids has not been well-defined. EVs are lipid bilayer delimited particles release by most cell types, can be found in biological fluids, and carry proteins, nucleic acids and lipids hypothesized to mediate intercellular communication by transfer of their cargo to recipient cells^16^. In the context of neurodegeneration, EVs can propagate pathogenic molecules, including Tau filaments^17–20^. In contrast, NVEPs are non-membranous particles shown to play a role in disease pathogenesis^21,12^ and carry proteins associated with various diseases, such as proprotein convertase subtilisin/kexin type 9 in cardiovascular diseases or amyloid precursor protein in AD^13,22,23^. NVEPs, have been classified into two subtypes of particles, exomeres and supermeres^22–24^. Like EVs, exomeres and supermeres may also contain functional cargo that can serve both as disease biomarkers and drive signaling and metabolic changes in recipient cells^12,21,25^. However, these subpopulations have not been widely studied in the CNS at the cell-type specific level as potential biomarkers of neurodegeneration. Therefore, EVs and specifically NVEPs represent an underexplored source of novel biomarkers of neuroinflammation. However, as these particles contain distinct molecular cargoes, methods to separate and characterize them in relevant biofluids like plasma need to be established to translate the pre-clinical findings to clinically relevant biomarkers.

Here, we systematically isolated and characterized the molecular and biophysical properties of microglia-derived EVs and NVEPs under pro-inflammatory or anti-inflammatory conditions. We profiled their small RNA cargo by sequencing and validated this signature in human iPSC-derived microglia. We further investigated the features of EVs and NVEPs derived from mouse plasma and brain tissue in an engineered mouse model for tracking microglia-specific EVs in the context of neuroinflammation. We found different glycosylation patterns on EVs and NVEPs under different activation states, which has only been previously identified in cancer-derived EVs and exomeres^26^. Our data show that microglia-derived EVs and NVEPs, along with their RNA cargo, reflect key aspects of microglial cellular states, and appear to be linked to neurodegenerative phenotypes in neurological disease. Finally, we demonstrated the use of immunoaffinity methods and single particle imaging in plasma as the optimal method to assay these distinct particles in isolation. Importantly, these findings provide support for further development of these markers for validation in clinical samples.

## Results

### Biophysical and molecular characterization of microglia-derived extracellular vesicles and non-vesicular extracellular particles

EVs and NVEPs were isolated from HMC3 cell conditioned media as previously described^27^ with some modifications (Fig. 1a and Extended Fig. 1). To characterize the size distribution, isolated EVs (large and small) and NVEPs (exomeres and supermeres) were imaged by transmission electron microscopy (TEM), which revealed distinct size differences, with NVEPs noted to be smaller than sEVs as previously reported (Fig.1b). Given their small size and inadequate resolution on TEM, we further characterized the size, morphology and mechanical properties of exomeres and supermeres using high-resolution atomic force microscopy (AFM). AFM imaging of both exomeres and supermeres allowed direct visualization of their topographical height profiles (Fig. 1c). The height and diameter measurements showed a height of 7.28 ± 0.08 nm for exomeres and 5.66 ± 0.03 nm (mean ± SEM) for supermeres. Similarly, diameters for exomeres and supermeres were found to be 26.6 ± 0.3 nm and 18.4 ± 0.2 nm (mean ± SEM), respectively. TEM (Fig. 1b) and protein quantification (Fig. 1d) analyses revealed a higher abundance of supermeres and exomeres in these preparations compared to the EV fractions (lEVs: 85.42 ± 47.35 µg/mL; sEVs: 3505 ± 459.1 µg/mL; exomeres: 10167 ± 2017 µg/mL; supermeres: 11165 ± 1918 µg/mL). Finally, EVs and NVEPs were further characterized by immunoblotting (Fig. 1e). We detected the presence of canonical EV markers (membrane EV markers: CD63, CD9, CD81, flotlin-1; and cargo EV marker: syntenin-1) and NVEP-associated markers (TGFbi, AGO2, ACE2, and GAPDH). Interestingly, low levels of tetraspanins were detected in the exomeres, suggesting either co-isolation of EV components in the exomeres fraction or presence of tetraspanins in NVEPs. Using nanoflow cytometry we analyzed typical surface EV markers (CD9, CD81, and CD63) in sEVs, exomeres, and supermeres fractions. Consistent with the immunoblotting data, these markers were present in both EVs and exomeres (positive population shown in red as P1) and rarely detected on supermeres (Fig. 1f).

**Figure 1:**
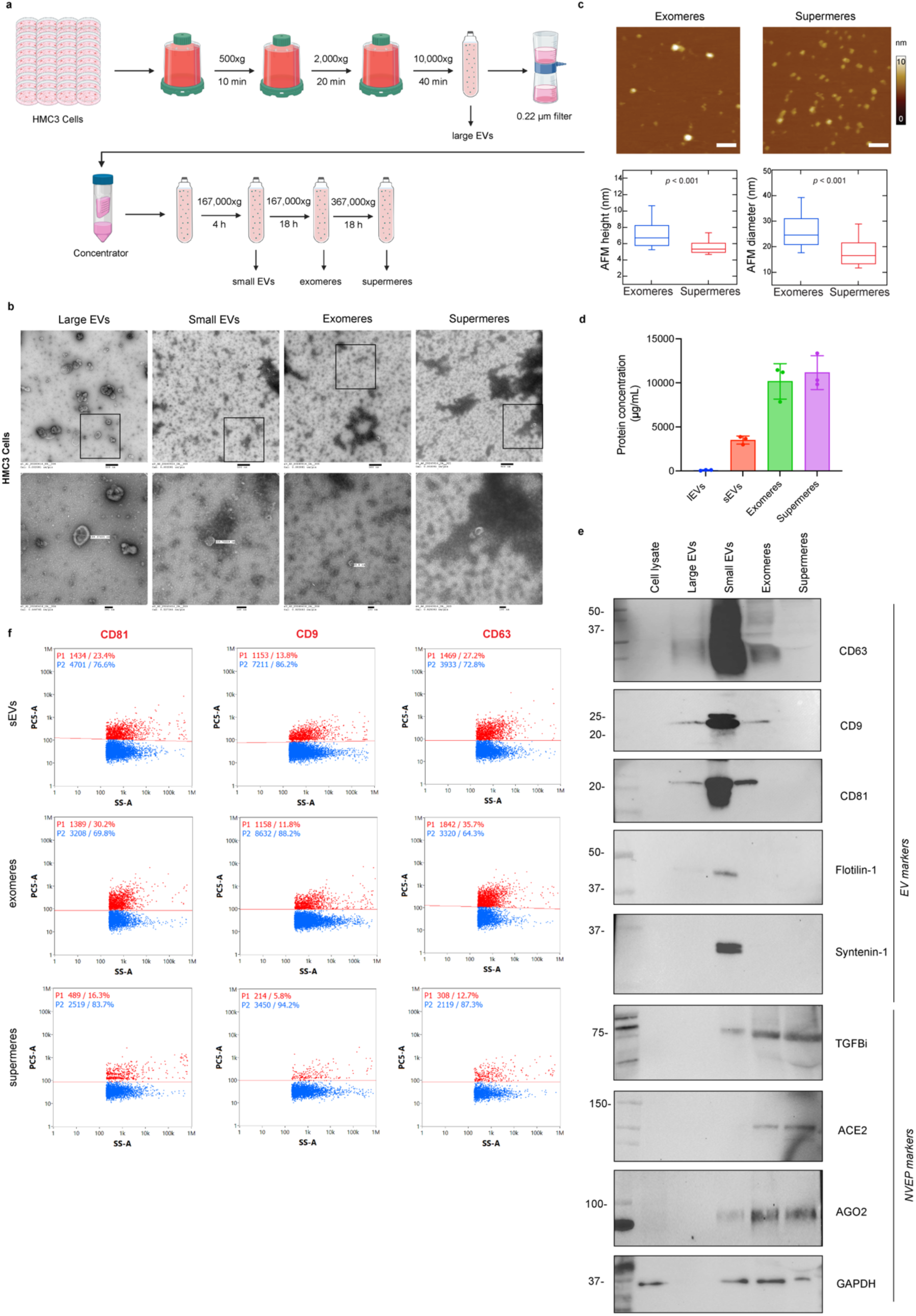
Characterization of microglia-derived extracellular vesicles and non-vesicular extracellular particles. **a**. Schematic representation of the EVs and NVEPs isolation protocol. **b**. Representative transmission electron microscopy images of isolated EVs and NVEPs. **c**. Representative tapping mode atomic force microscopy (AFM) images of isolated exomeres and supermeres derived from microglial cells. Scale bars, 100 nm. Exomeres and supermeres height and diameters measured by AFM. Box and whisker plot of height and diameter of Exomeres (N = 572) and Supermeres (N = 823). The center lines of the box plot mark the median, the box limits indicate the 25th and 75th percentiles and the lower and upper limits of whiskers are at the 9th and 91st percentiles, respectively. The diamond point inside the box indicates the respective means. For the statistical significance of data, a two-tailed Student’s t-test was performed, which gave p <0.001 for both diameter and height data. **d**. Protein concentration measured by BCA assay. Data represented as mean ± SD. **e**. Immunoblot of selected EV and NVEP markers in microglia cell lysates, large EVs, small EVs, exomeres and supermeres. **f**. Nanoflow cytometry of selected markers CD9, CD81 and CD63 (positive in red and negative populations in blue) for microglia-derived small EVs, exomeres and supermeres.

### Characterization of microglia-derived extracellular vesicles and non-vesicular extracellular particles under pro-inflammatory and anti-inflammatory conditions

HMC3 microglial cells were cultured under pro-inflammatory (LPS, 100 ng/mL) or anti-inflammatory (IL-10, 50 ng/mL) conditions for 48 h to model distinct activation states. Conditioned media were subsequently collected for the isolation of EVs and NVEPs. Under pro-inflammatory conditions microglia showed upregulation of M1 markers (IL-6) and downregulation of M2 markers (CD206) (Extended Fig. 2a). Compared to untreated controls or the anti-inflammatory group, LPS treatment induced a significant increase in cell size, with microglia displaying enlarged mitochondria as observed by immunofluorescence (Fig. 2a). Lectin staining with Sambucus nigra agglutinin (SNA), which specifically binds α2,6-linked sialic acid residues, showed comparable levels of cell surface glycosylation across different microglial activation states (Fig. 2b). This indicates that microglial activation does not markedly alter α2,6-sialylation at the plasma membrane. By first assessing α2,6-sialylation directly on microglial cells, we established the baseline cellular glycosylation profile, which served as a reference for subsequent analyses of secreted extracellular particles. Protein quantification by BCA assay revealed that exomeres and supermeres contained higher amounts of protein compared to other fractions (Extended Fig. 2b). EVs and NVEPs were characterized by immunoblotting (Fig. 2c). Canonical EV markers were detected, including membrane-associated proteins tetraspanins CD63, CD9, CD81 (TSPAN) and the cargo marker Alix, along with NVEP-associated markers LDHA, TGFBi, AGO2 and SCTH among the different microglia states. Multiple resistive pulse sensing analysis revealed that large EVs measured approximately 200-400 nm (Fig. 2d), while small EVs measured around 50-100 nm (Fig. 2f) consistent with previous descriptions. No differences in particle concentration for small and large EVs were observed between groups (Fig. 2e and 2g). TSPANs are abundant on most EVs^28,29^ but have low abundance in NVEPs^12,21,22^. However, due to the characteristics of the ultrafiltration/differential ultracentrifugation isolation method, TSPANs were also detected in exomeres fractions and, to a lesser extent, in supermeres fractions (Extended Fig. 2c).

**Figure 2:**
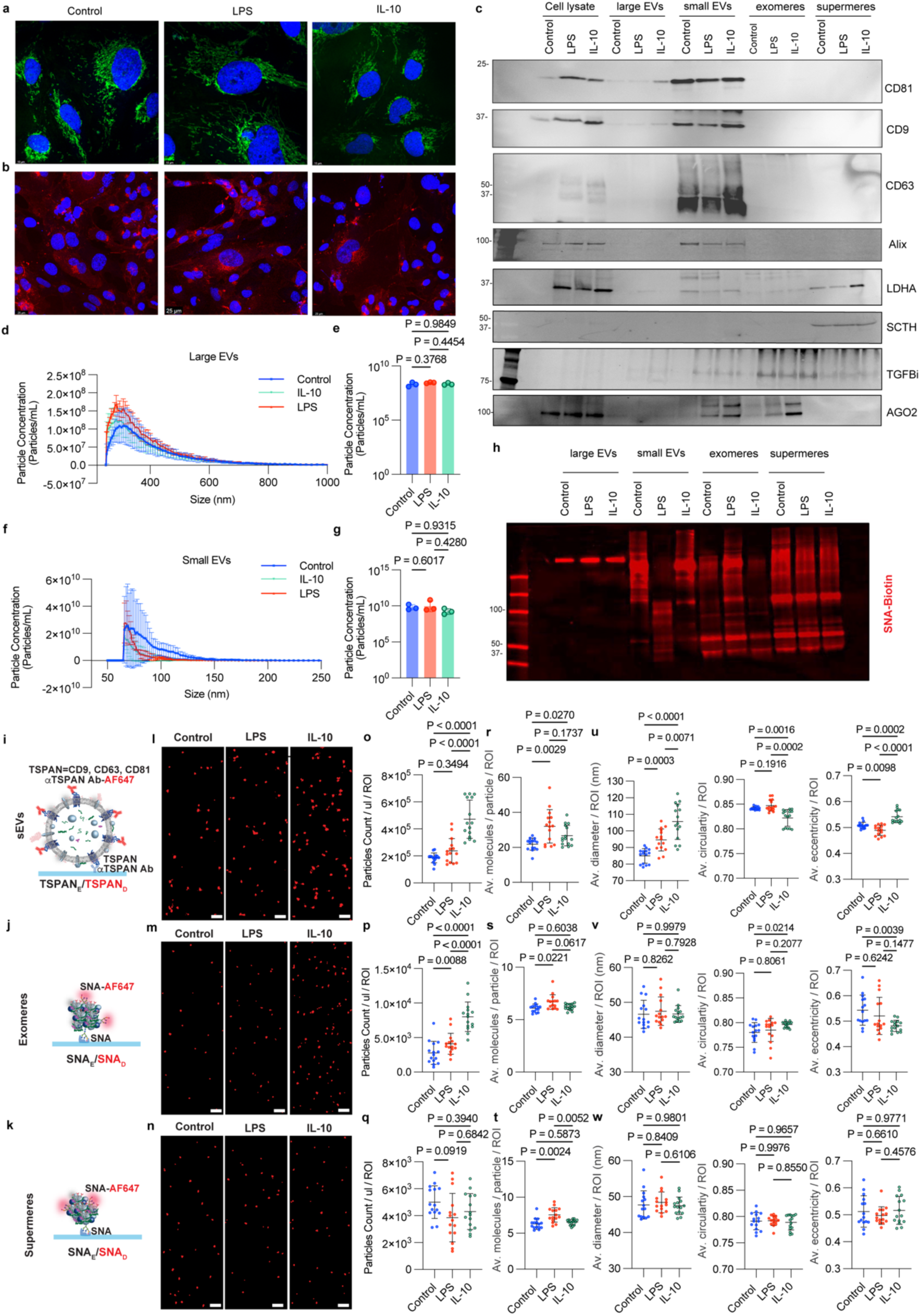
Distinct signatures of state-specific microglia-derived extracellular vesicles and non-vesicular extracellular particles. **a.** Representative images of HMC3 cells under control, pro-inflammatory or anti-inflammatory conditions stained for mitochondria marker COXIV. Nuclei were labelled by DAPI. Scale bar: 10 μm. **b.** Representative images of the cellular distribution of α2,6-sialylation in HMC3 cells under control, pro-inflammatory or anti-inflammatory conditions. α2,6-sialylation characterized by the staining of biotin-labeled *Sambucus nigra agglutinin* (SNA). Nuclei were labelled by DAPI. Scale bar: 25 μm. **c.** Immunoblot of selected EV and NVEP markers in HMC3 microglia cell lysates, large EVs, small EVs, exomeres and supermeres. **d.** Microfluidic resistive pulse sensing (MRPS) data of isolated large EVs. Data represented as Mean ± SD (*n*=3 samples per group). **e.** Comparison of large EVs concentration across samples measured by MRPS (*n*=3 samples per group). **f.** Microfluidic resistive pulse sensing (MRPS) data of isolated small EVs. Data represented as Mean ± SD (*n*=3 samples per group). P-values were calculated by ANOVA. **g.** Comparison of small EVs concentration across samples measured by MRPS. Data represented as mean ± SD. (*n*=3 samples per group). P-values were calculated by ANOVA. **h.** Representative images of SNA blot analysis of cell lysates, EVs (large and small EVs) and NVEPs (exomeres and supermeres). **i-j.** Schematic representation of the detection and capture of sEVs, exomeres and supermeres respectively. **l-m.** Representative images of EVs, exomeres and supermeres under control, pro-inflammatory or anti-inflammatory conditions. Scale bar 500 nm. **o-q.** Number of detected particles for microglia isolated EVs and NVEPs. Data was normalized per 1 μl of purified particles. **r-t.** Number of detected cargo molecules per particle (TSPANs for sEVs and SNA detected glycans for exomeres/supermeres). **u-w**. Quantification of morphology: size (left), circularity (middle), eccentricity (right). Averages ± SEM are shown (n=3 independent fractions, each imaged with 5 ROI).

NVEPs display distinct N-glycosylation patterns^21,26^. For example, supermeres and exomeres have unique staining pattern with SNA lectin that recognizes α-2,6-linked sialic acid and SNA has been used to isolate/precipitate NVEPs^21^. We have confirmed unique SNA staining patterns on NVEPs despite the absence of detectable changes at the cellular level (Fig. 2h). Based on these observations, we developed a method to characterize sEVs, exomeres, and supermeres using Single Extracellular VEsicle Nanoscopy (SEVEN)^30^. This approach combines affinity isolation with fluorescence-based super-resolution microscopy to provide information on EV/NVEP morphology and molecular cargo content. We detected EVs via TSPAN capture/detection and NVEPs (exomeres and supermeres) via SNA capture/detection (Fig. 2i–k). Using this approach, we clearly visualized defined structures of isolated EVs and NVEPs with excellent signal-to-noise resolution; minimal signal was observed with buffer controls (Table S1). Raw images (Fig. 2l–n) point to distinct sizes and morphology between EVs and exomeres/supermeres. Under anti-inflammatory conditions, we observed a significant increase in the number of secreted sEVs and exomeres, whereas supermeres secretion was unaffected (Fig. 2o-q, Table S1). Pro-inflammatory conditions induced a modest but significant increase in exomeres secretion, with no effect on sEVs or supermeres. In addition, for each detected particle, we obtained the number of detected cargo molecules (TSPANs for EVs and SNA detected glycans for NVEPs). Pro-inflammatory conditions significantly increased molecular cargo content in sEVs, exomeres, and supermeres, whereas anti-inflammatory conditions induced a modest but significant increase only in sEVs cargo (Fig. 2r-t). Morphological quantification of diameter and shape (circularity measures how close particle approximates a perfect circle while eccentricity measures particle elongation) further revealed distinct features (Fig. 2u-w). As expected, compared to EVs, NVEPs were significantly smaller, had lower size heterogeneity, and were less circular. Pro-inflammatory conditions modestly altered sEVs morphology by significantly increasing size and reducing elongation, without affecting exomeres or supermeres. By contrast, anti-inflammatory conditions profoundly altered sEVs morphology, significantly increasing size (and size heterogeneity), reducing circularity, and increasing elongation. Exomeres showed increased circularity and reduced elongation under anti-inflammatory conditions, while supermeres morphology remained unchanged. Descriptive statistics for EV and NVEP properties under three conditions are shown in Table S2 (average value per region of interest, ROI) and Table S3 (average values for all detected EVs/NVEPs) and revel their distinct signatures.

### EVs and NVEPs exhibit distinct small RNA signatures under pro-inflammatory and anti-inflammatory conditions

Total RNA was isolated from three different preparations of control, pro-inflammatory, or anti-inflammatory microglia-derived sEVs and NVEP (exomeres and supermeres together), and RNA integrity was analyzed using a bioanalyzer. The bioanalyzer gel-image showed that EVs are enriched in both long and small RNA species while NVEPs were predominantly enriched in small RNAs (Fig. 3a and b). EVs from LPS-treated HMC3 cells contained higher total RNA levels compared to control EVs (control: 901.7 ± 257.1 ng; LPS: 1310 ± 36.06 ng; IL-10: 1015 ± 72.15 ng) (Fig. 3c). No significant differences in total RNA concentration were observed among NVEPs across conditions (control: 675 ± 602.2 ng; LPS: 546 ± 250.7 ng; IL-10: 337.3 ± 212.9 ng) (Fig. 3d). We then carried out RNA sequencing on the three microglial states (Control, LPS or IL-10, *n*=3 samples per group) of each EVs and NVEPs group. Analysis of RNA sequencing reads from the isolated fractions revealed a diverse composition of RNA biotypes. A substantial proportion of the reads mapped to tRNAs and YRNAs, indicating that these small non-coding RNAs are highly abundant in both EVs and NVEPs (Fig. 3e). Further analysis of unique transcripts showed that miRNAs were the most diverse RNA group in both EVs and NVEPs (Fig. 3f). Distinct miRNA signatures were noted for each of these states in the sEVs as well as NVEPs (Fig. 3g and 3h). Among the miRNAs differentially expressed between microglial states in both EVs and NVEPs hsa-miR-146a-5p was upregulated upon LPS treatment in both EVs and NVEPs (Fig. 3i and 3j). In EVs hsa-miR-10399-5p was also upregulated upon LPS treatment, whereas NVEPs had fewer differentially expressed miRNAs overall (Table 3 and 4). Volcano plot analyses further complemented the differences in EV-miRNA signatures revealed by the heatmaps. In EVs, several miRNAs were significantly upregulated in control versus anti-inflammatory and control versus pro-inflammatory comparisons, with distinct subsets unique to each condition (Fig. 3k and 3l). NVEPs displayed fewer significant changes (Fig. 3m and 3n), but still exhibited state specific alterations, suggesting that small RNA cargo sorting into EVs and NVEPs is differentially regulated under inflammatory versus anti-inflammatory states. Comparison of EVs and NVEPs small RNAs revealed unique genes in EVs and in NVEPs (Extended Data Fig. 3c).

**Figure 3:**
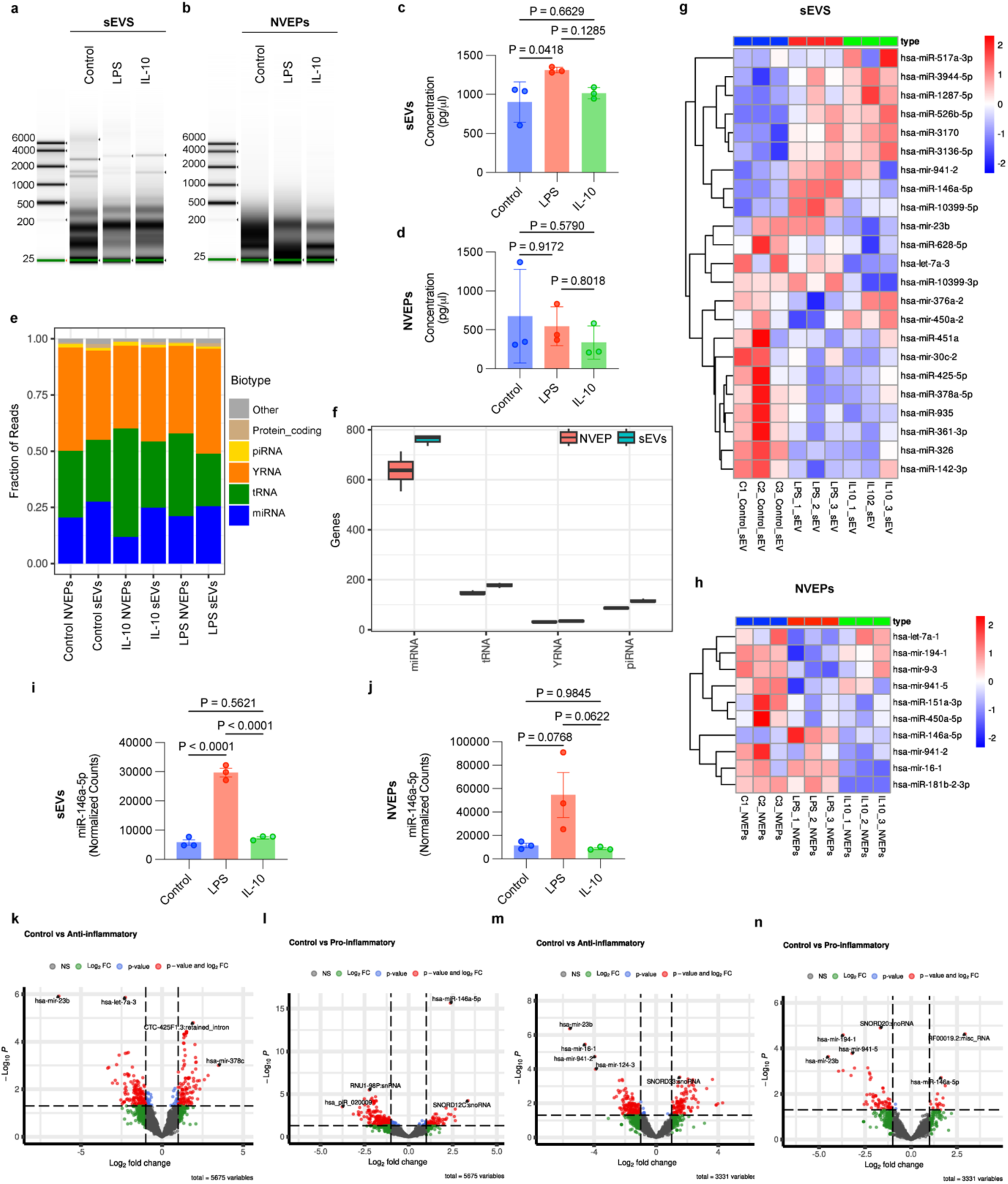
Characterization of small RNA composition in control, pro-inflammatory or anti-inflammatory microglia-derived extracellular vesicles and non-vesicular extracellular nanoparticles. **a.** Bioanalyzer image of RNA quality extracted from sEV samples (*n*=3 samples per group). **b.** Bioanalyzer image of RNA quality extracted from NVEP samples (*n*=3 samples per group). **c**. Comparison of total RNA yield extracted from EV samples (*n*=3 samples per group, *p-value =0.0418*). P-values were calculated by ANOVA with Tukey’s HSD test. **d**. Comparison of total RNA yield extracted from NVEP samples (*n*=3 samples per group). **e**. Distribution average of RNA biotypes on sEVs and NVEPs samples (*n*=3 samples per group). **f**. Box plots of unique genes detected in sEVs and NVEPs (*n*=3 samples per group). **g.** Heatmaps showing differentially expressed miRNAs in sEVs (*n*=3 samples per group). **h**. Heatmaps showing differentially expressed miRNAs in NVEPs (*n*=3 samples per group). **i**. Box plots showing the expression of miR-146a-5p in EVs with respective p-values using Mann Whitney t-test (*n*=3 samples per group). Mean ± SD. **j**. Box plots showing the expression of miR-146a-5p in NVEPs (*n*=3 samples per group). Mean ± SD. Volcano plots showing differential expression of small RNAs in extracellular vesicles (EVs) **k**. Control vs. anti-inflammatory conditions. **l.** Control vs. pro-inflammatory conditions. Volcano plots showing differential expression of small RNAs in non-vesicular extracellular particles (NVEPs). **m**. Control vs. anti-inflammatory conditions. **n.** Control vs. pro-inflammatory conditions. The x-axis represents log2fold change and the y-axis represents –log10 adjusted p-value. Dots highlight small RNAs significantly dysregulated (red) under each condition compared to control, with non-significant RNAs shown in black.

**Table 1:**
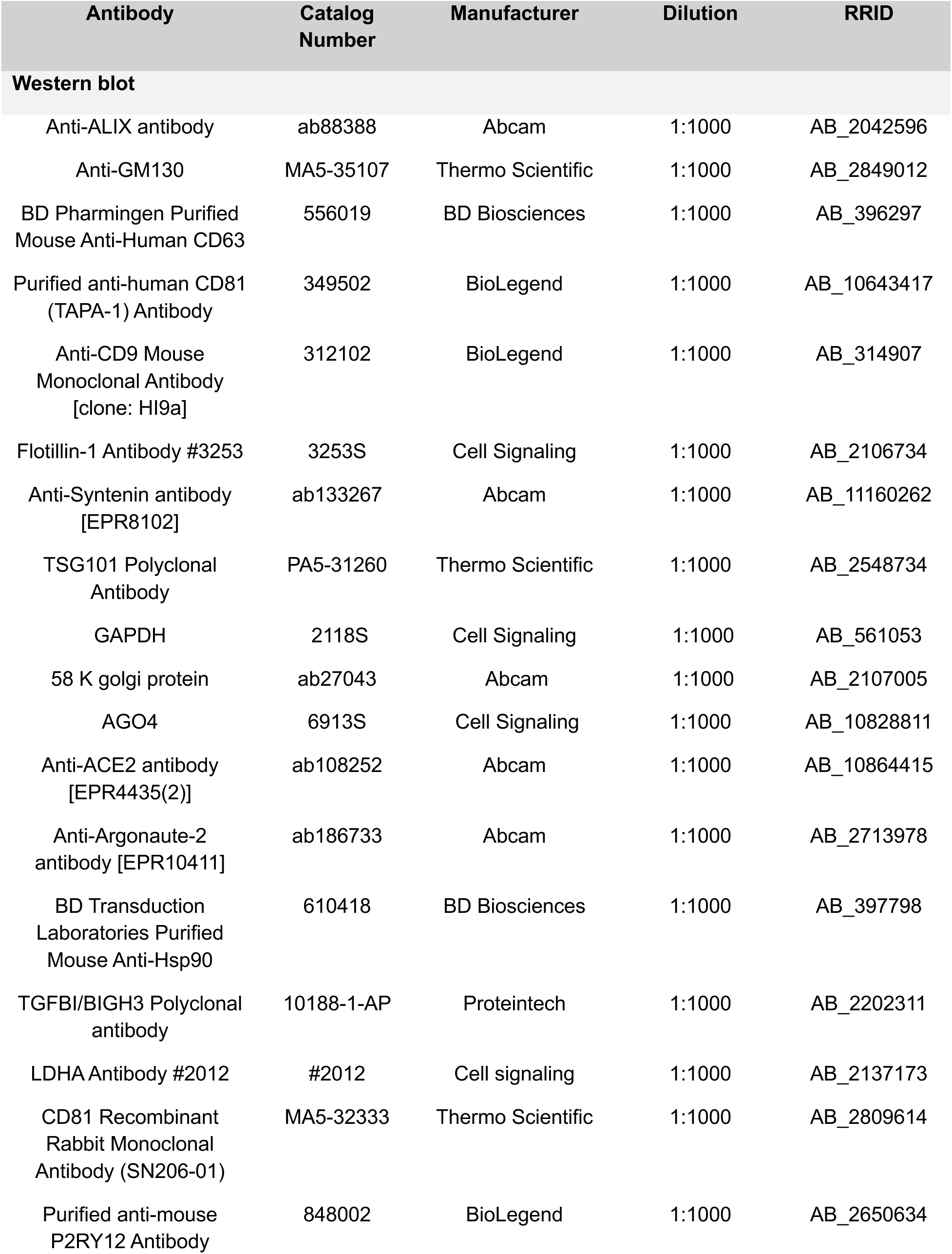
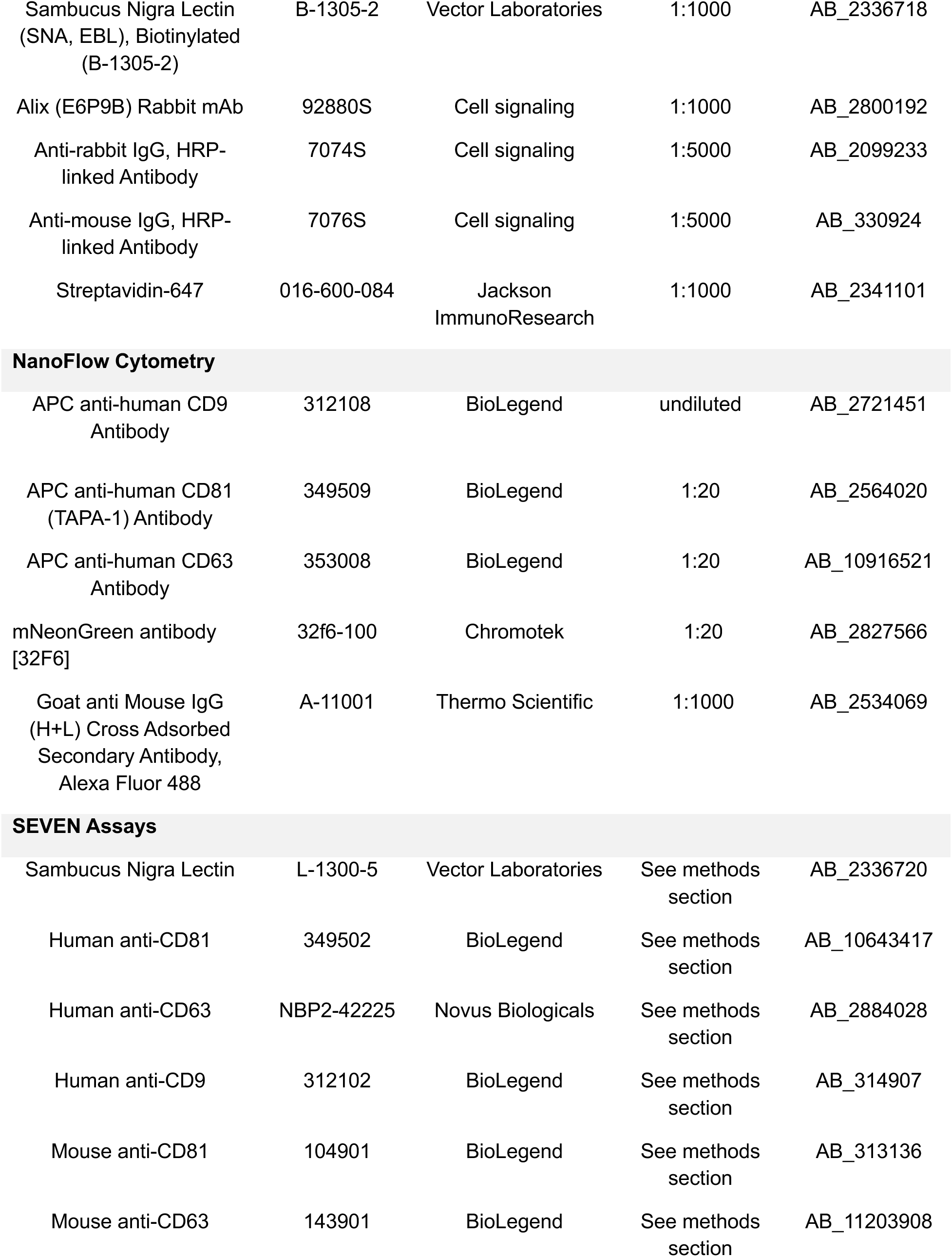
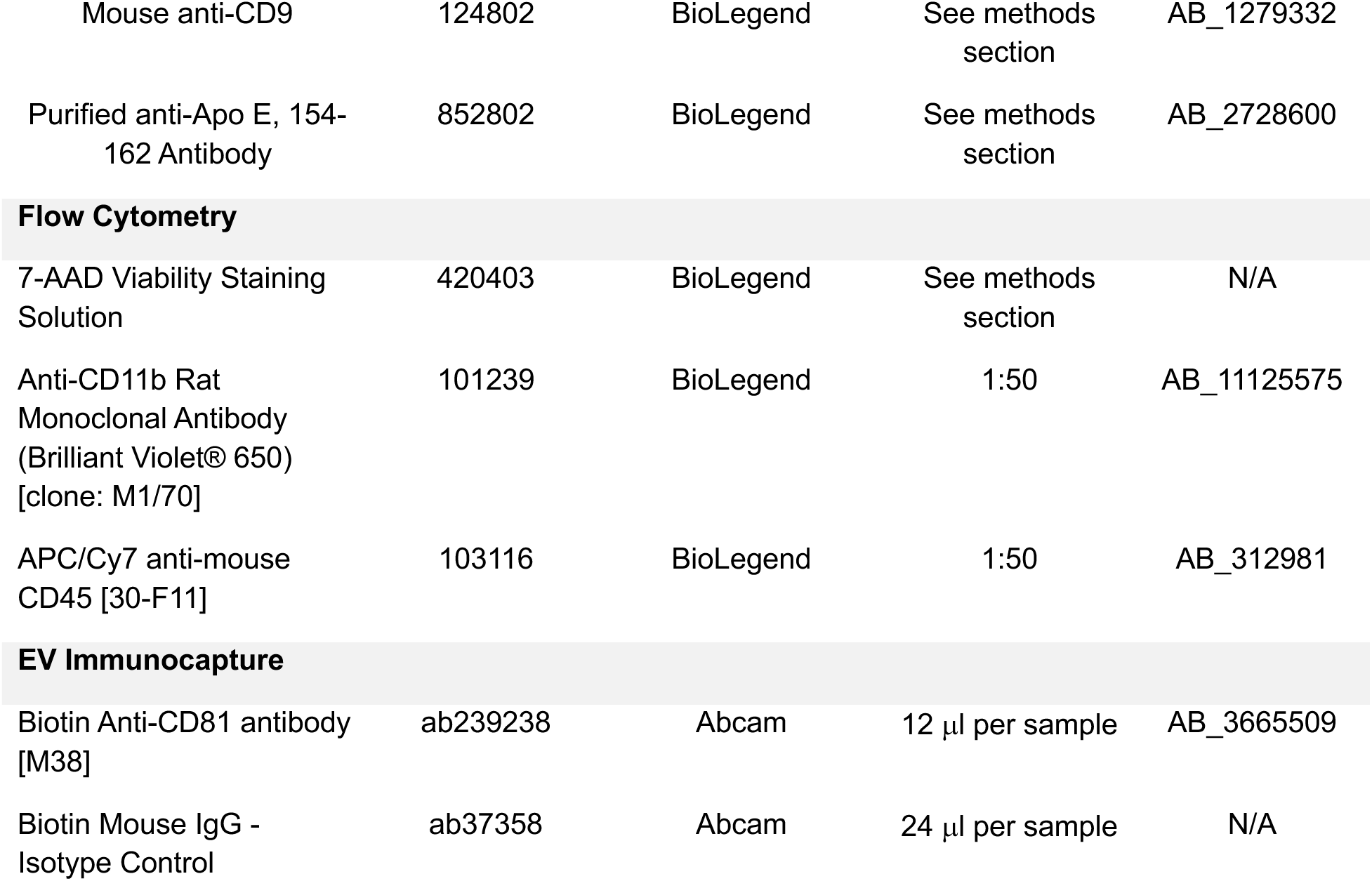
List of Antibodies.

| Antibody | Catalog Number | Manufacturer | Dilution | RRID |
| --- | --- | --- | --- | --- |
| **Western blot** |  |  |  |  |
| Anti-ALIX antibody | ab88388 | Abcam | 1:1000 | AB_2042596 |
| Anti-GM130 | MA5-35107 | Thermo Scientific | 1:1000 | AB_2849012 |
| BD Pharmingen Purified Mouse Anti-Human CD63 | 556019 | BD Biosciences | 1:1000 | AB_396297 |
| Purified anti-human CD81 (TAPA-1) Antibody | 349502 | BioLegend | 1:1000 | AB_10643417 |
| Anti-CD9 Mouse Monoclonal Antibody [clone: HI9a] | 312102 | BioLegend | 1:1000 | AB_314907 |
| Flotillin-1 Antibody #3253 | 3253S | Cell Signaling | 1:1000 | AB_2106734 |
| Anti-Syntenin antibody [EPR8102] | ab133267 | Abcam | 1:1000 | AB_11160262 |
| TSG101 Polyclonal Antibody | PA5-31260 | Thermo Scientific | 1:1000 | AB_2548734 |
| GAPDH | 2118S | Cell Signaling | 1:1000 | AB_561053 |
| 58 K golgi protein | ab27043 | Abcam | 1:1000 | AB_2107005 |
| AGO4 | 6913S | Cell Signaling | 1:1000 | AB_10828811 |
| Anti-ACE2 antibody [EPR4435(2)] | ab108252 | Abcam | 1:1000 | AB_10864415 |
| Anti-Argonaute-2 antibody [EPR10411] | ab186733 | Abcam | 1:1000 | AB_2713978 |
| BD Transduction Laboratories Purified Mouse Anti-Hsp90 | 610418 | BD Biosciences | 1:1000 | AB_397798 |
| TGFBI/BIGH3 Polyclonal antibody | 10188-1-AP | Proteintech | 1:1000 | AB_2202311 |
| LDHA Antibody #2012 | #2012 | Cell signaling | 1:1000 | AB_2137173 |
| CD81 Recombinant Rabbit Monoclonal Antibody (SN206-01) | MA5-32333 | Thermo Scientific | 1:1000 | AB_2809614 |
| Purified anti-mouse P2RY12 Antibody | 848002 | BioLegend | 1:1000 | AB_2650634 |
| Sambucus Nigra Lectin (SNA, EBL), Biotinylated (B-1305-2) | B-1305-2 | Vector Laboratories | 1:1000 | AB_2336718 |
| Alix (E6P9B) Rabbit mAb | 92880S | Cell signaling | 1:1000 | AB_2800192 |
| Anti-rabbit IgG, HRP-linked Antibody | 7074S | Cell signaling | 1:5000 | AB_2099233 |
| Anti-mouse IgG, HRP-linked Antibody | 7076S | Cell signaling | 1:5000 | AB_330924 |
| Streptavidin-647 | 016-600-084 | Jackson ImmunoResearch | 1:1000 | AB_2341101 |
| NanoFlow Cytometry |  |  |  |  |
| APC anti-human CD9 Antibody | 312108 | BioLegend | undiluted | AB_2721451 |
| APC anti-human CD81 (TAPA-1) Antibody | 349509 | BioLegend | 1:20 | AB_2564020 |
| APC anti-human CD63 Antibody | 353008 | BioLegend | 1:20 | AB_10916521 |
| mNeonGreen antibody [32F6] | 32f6-100 | Chromotek | 1:20 | AB_2827566 |
| Goat anti Mouse IgG (H+L) Cross Adsorbed Secondary Antibody, Alexa Fluor 488 | A-11001 | Thermo Scientific | 1:1000 | AB_2534069 |
| SEVEN Assays |  |  |  |  |
| Sambucus Nigra Lectin | L-1300-5 | Vector Laboratories | See methods section | AB_2336720 |
| Human anti-CD81 | 349502 | BioLegend | See methods section | AB_10643417 |
| Human anti-CD63 | NBP2-42225 | Novus Biologicals | See methods section | AB_2884028 |
| Human anti-CD9 | 312102 | BioLegend | See methods section | AB_314907 |
| Mouse anti-CD81 | 104901 | BioLegend | See methods section | AB_313136 |
| Mouse anti-CD63 | 143901 | BioLegend | See methods section | AB_11203908 |
| Mouse anti-CD9 | 124802 | BioLegend | See methods section | AB_1279332 |
| Purified anti-Apo E, 154-162 Antibody | 852802 | BioLegend | See methods section | AB_2728600 |
| **Flow Cytometry** |  |  |  |  |
| 7-AAD Viability Staining Solution | 420403 | BioLegend | See methods section | N/A |
| Anti-CD11b Rat Monoclonal Antibody (Brilliant Violet® 650) [clone: M1/70] | 101239 | BioLegend | 1:50 | AB_11125575 |
| APC/Cy7 anti-mouse CD45 [30-F11] | 103116 | BioLegend | 1:50 | AB_312981 |
| **EV Immunocapture** |  |  |  |  |
| Biotin Anti-CD81 antibody [M38] | ab239238 | Abcam | 12 µl per sample | AB_3665509 |
| Biotin Mouse IgG - Isotype Control | ab37358 | Abcam | 24 µl per sample | N/A |

**Table 2:**
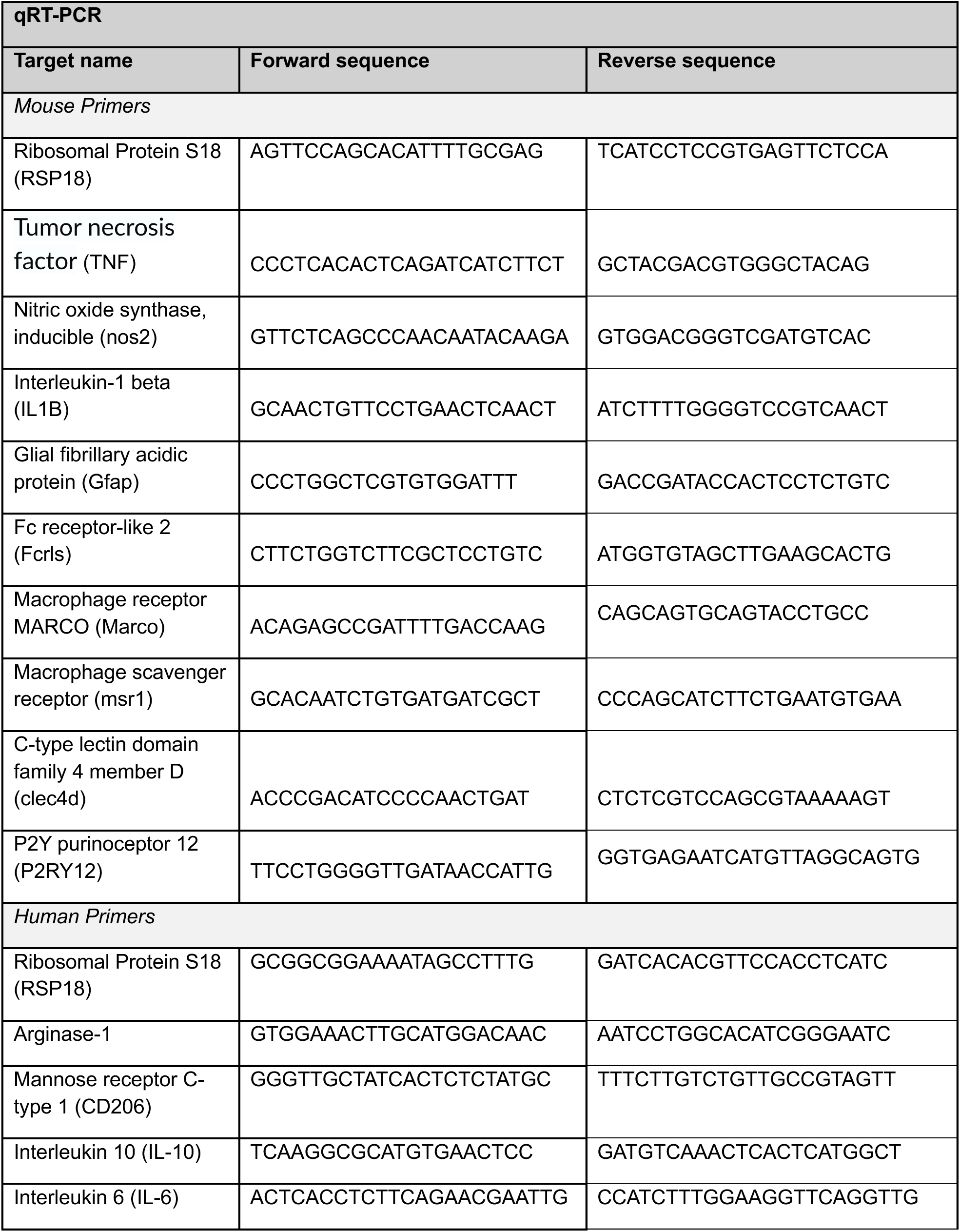
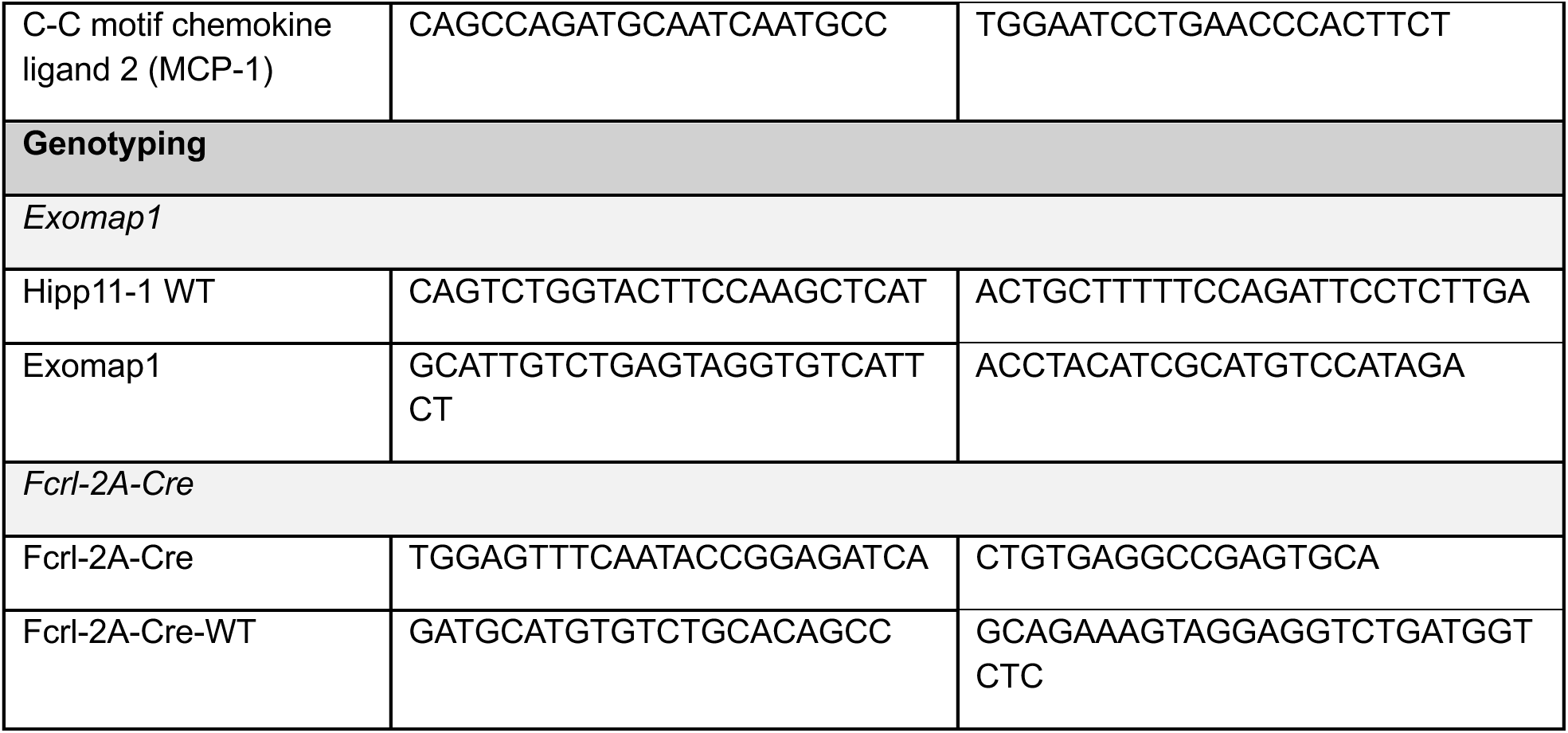
List of Primers.

**Table 3:**
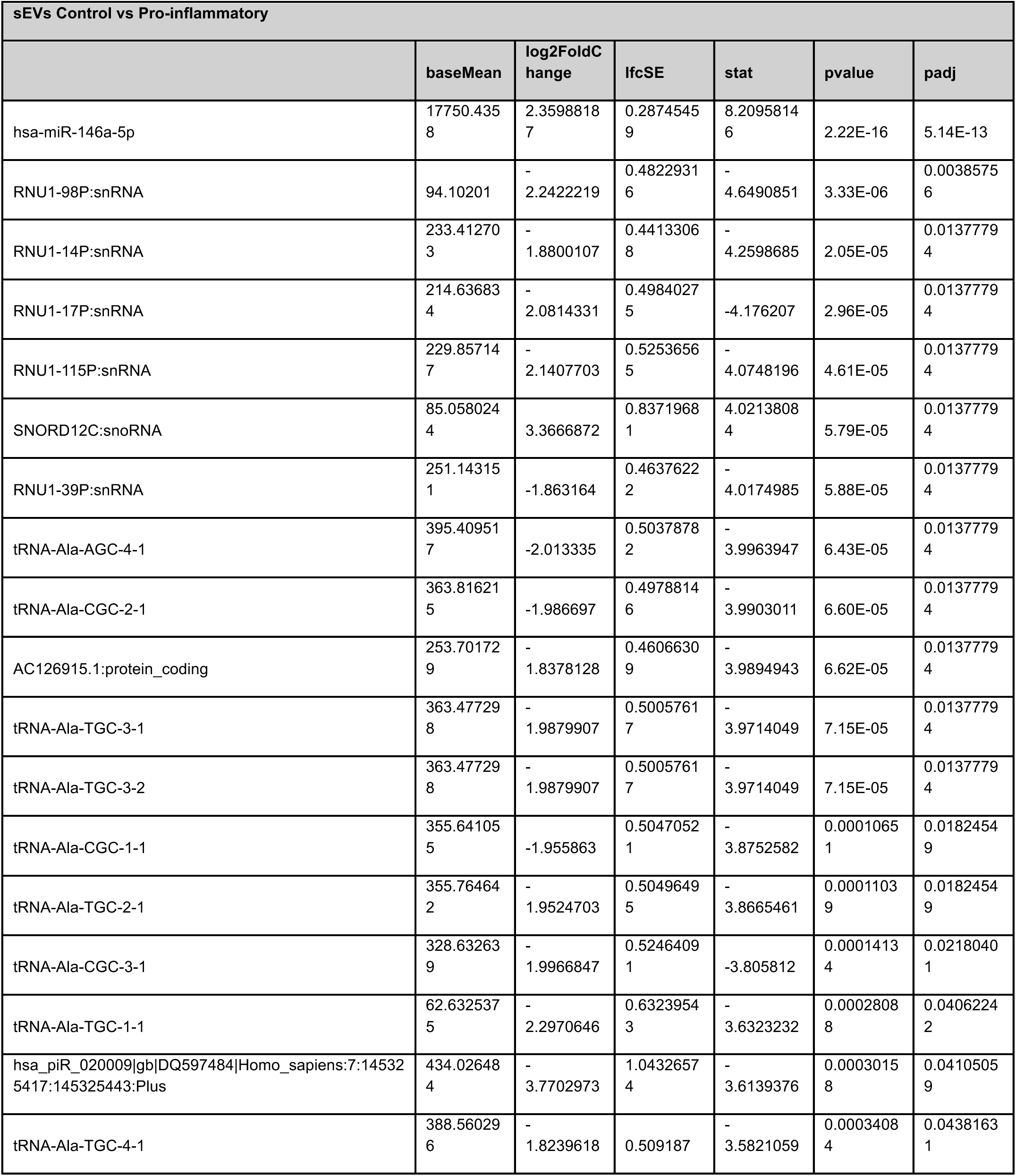
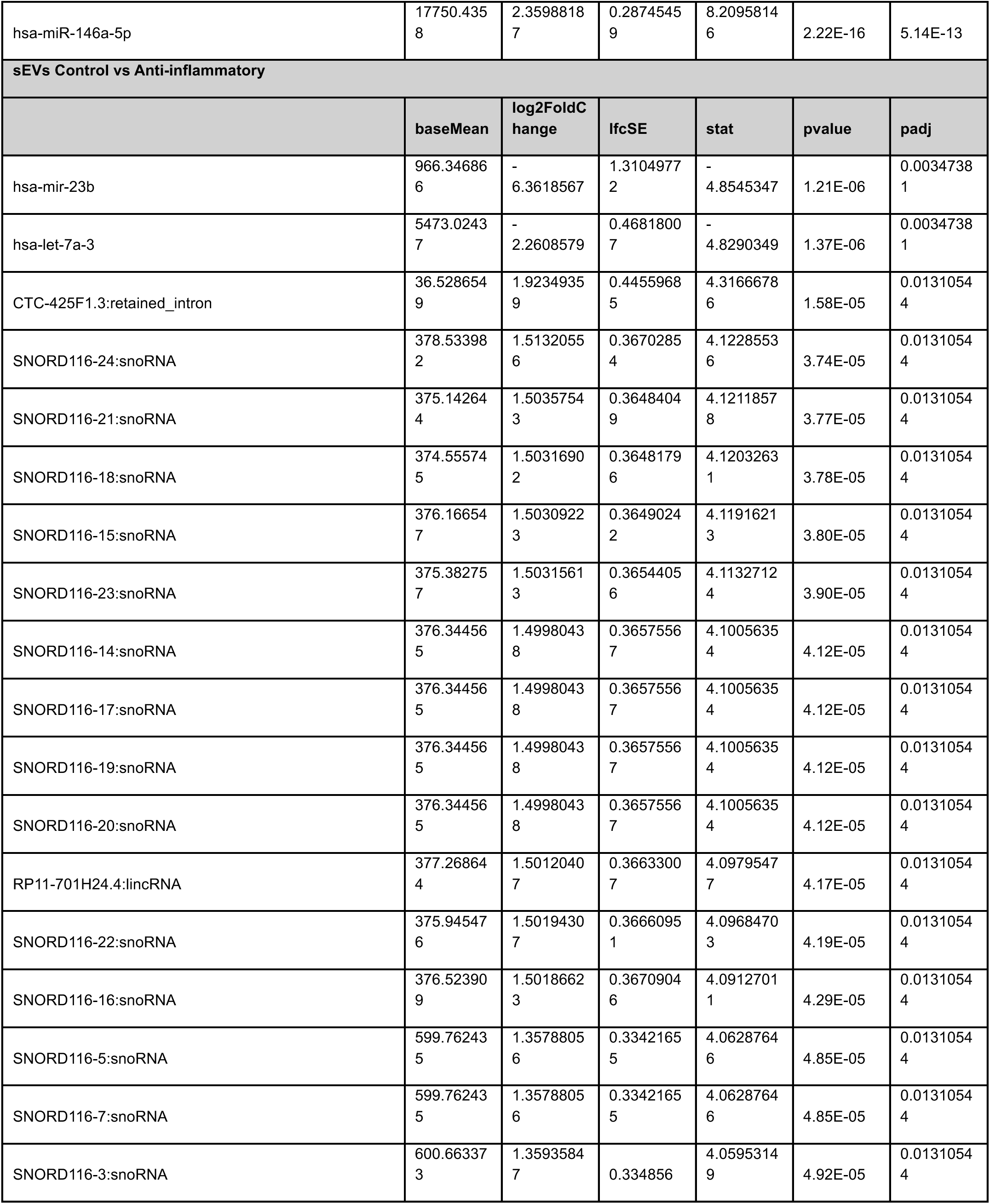
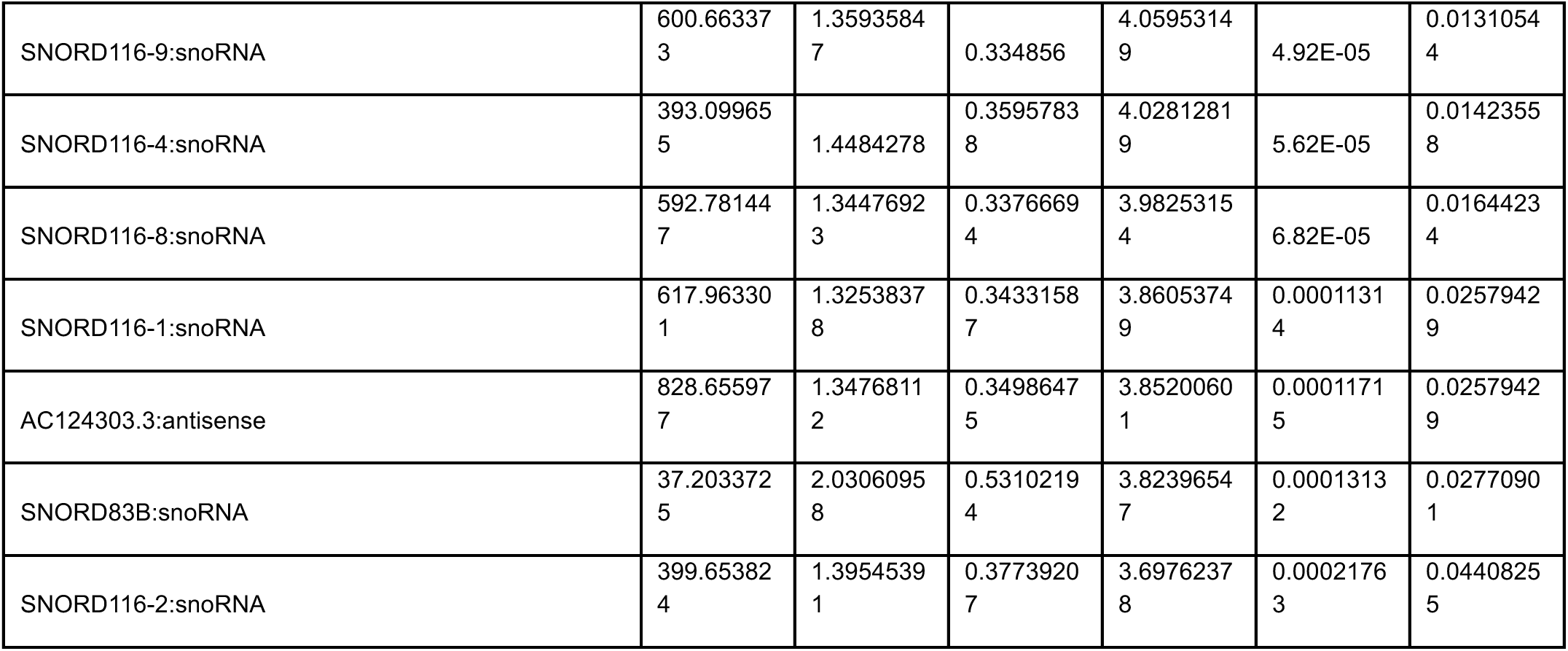
Differentially expressed small RNAs in HMC3 cell-derived EVs under pro-inflammatory and anti-inflammatory conditions.

**Table 4:** Differentially expressed small RNAs in HMC3 cell-derived NVEPs under pro-inflammatory and anti-inflammatory conditions.

| NVEPs Control vs Pro-inflammatory |  |  |  |  |  |  |
| --- | --- | --- | --- | --- | --- | --- |
|  | baseMean | log2FoldChange | lfcSE | stat | pvalue | padj |
| hsa-mir-194-1 | 22.9097383 | -4.350836 | 0.93381131 | -4.659224 | 3.17E-06 | 0.01194708 |
| hsa-miR-146a-5p | 20323.4622 | 2.24891883 | 0.55864606 | 4.02565952 | 5.68E-05 | 0.07274793 |
| hsa-mir-9-3 | 54.4771096 | -3.7453591 | 0.94270878 | -3.9729757 | 7.10E-05 | 0.07274793 |
| hsa-mir-941-5 | 381.70014 | -3.7379274 | 0.94569017 | -3.9525921 | 7.73E-05 | 0.07274793 |
| NVEPs Control vs Anti-inflammatory |  |  |  |  |  |  |
|  | baseMean | log2FoldChange | lfcSE | stat | pvalue | padj |
| hsa-mir-16-1 | 228110.272 | -4.621951 | 0.8716948 | -5.3022583 | 1.14E-07 | 0.00043052 |
| hsa-miR-181b-2-3p | 11.3543583 | -6.5488224 | 1.39324045 | -4.7004251 | 2.60E-06 | 0.00488606 |

### Human iPSC-derived iTF-Microglia recapitulate inflammatory signatures in cells, EVs, and NVEPs

Because HMC3 microglial cells may not fully capture the molecular and functional characteristics of adult human microglia, we next examined these phenotypes using human induced pluripotent stem cell (iPSC)-derived iTF-Microglia. These iPSC-derived microglia more closely resemble primary human microglia in gene expression, morphology, and function, providing a more physiologically relevant model for studying microglial states and functions^31^.During differentiation of the iPSCs into iTF-microglia, cells underwent morphological changes consistent with a microglial phenotype, including the transition from a rounded, progenitor-like shape to a ramified morphology with fine branching processes typical of resting microglia (Fig. 4a). Cells were stained for the microglial marker Iba1 (Fig. 4b), and transcriptional analyses showed downregulation of pluripotency genes (SOX2, NANOG, NODAL) (Fig. 4c) and upregulation of homeostatic microglial markers such as TMEM119, HEXB, and SELPLG (Fig. 4d). Conditioned media was collected, and sEVs and NVEPs were isolated and characterized by immunoblotting (Fig. 4e) and TSPAN ELISA (Extended Fig. 4a), which revealed the presence of tetraspanins on EVs and the NVEP-associated marker TGFBi. Nanoflow cytometry further demonstrated that the TSPAN CD63 was predominantly detected on sEVs (Fig. 4f). To mimic pro-inflammatory and anti-inflammatory microglial states, cells were treated with LPS (100 ng/mL) or IL-10 (50 ng/mL) for 48 hours. Cell enlargement, as well as mitochondrial fragmentation and enlargement, were observed by immunofluorescence (Fig. 4g). iTF-Microglia under pro-inflammatory conditions showed upregulation of M1 markers (IL-6 and MCP-1) and downregulation of M2 markers (Arginase-1, CD206, IL-10) (Fig. 4h). We further isolated total RNA cargo and analyzed it using a Bioanalyzer (Fig. 4i). Following isolation of RNAs from isolated EVs (Fig. 4i) we next assayed for the presence of our most promising EV-miRNA marker for inflammation miR-146a-5p, in cells, EVs, exomeres and supermeres. In cells, EVs and exomeres miR-146a-5p was upregulated under pro-inflammatory conditions, while it was not in supermeres (Fig. 4j). Consistent with our prior results, the upregulation was most notable in the sEVs.

**Figure 4:**
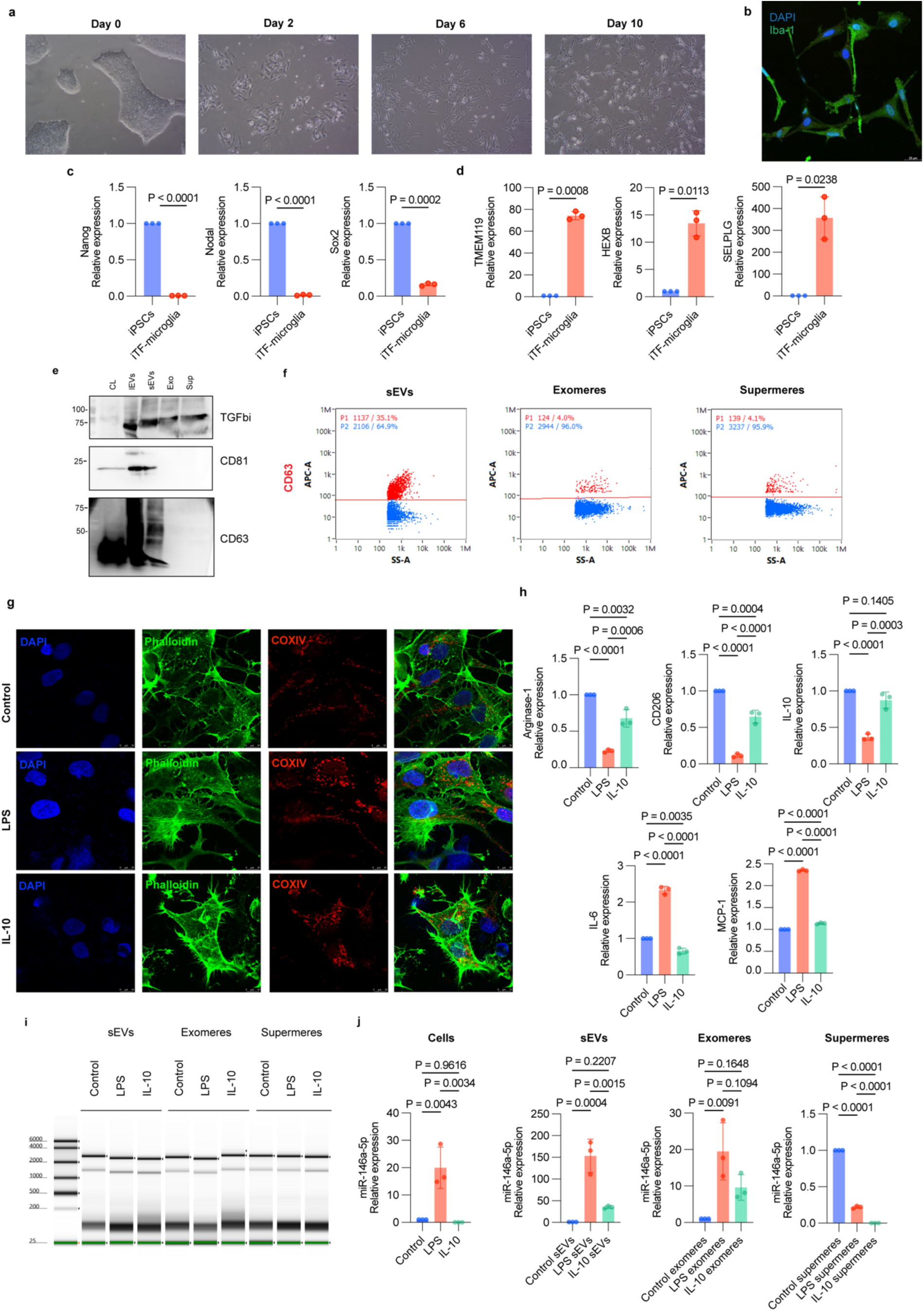
Differentiation and characterization of human iTF-microglia and their extracellular vesicles (EVs) and non-vesicular extracellular particles (NVEPs). **a.** Representative images during the differentiation process for generating iTF-Microglia on the indicated days. **b.** immunofluorescence image of iTF-Microglia on Day 15 of differentiation stained for microglia marker IBA1. Nuclei were labelled by DAPI. Scale bar: 25 μm. **c.** Quantitative RT-PCR analysis of iPSC genes Nanog, Nodal and Sox2 mRNA expression in iPSC and differentiated iTF-microglia. Rsp18 was used as an endogenous control to normalize gene expression levels. Relative mRNA abundance in iPSCs to iTF-microglia is reported. Results represent mean ± SD (Representative experiment with *n* = 3 technical replicates). **d.** Quantitative RT-PCR analysis of associated microglia markers TMEM119, HEXB and SELPLG mRNA expression in iPSC and differentiated iTF-microglia. Rsp18 was used as an endogenous control to normalize gene expression levels. Relative mRNA abundance in iPSCs to iTF-microglia is reported. Results represent mean ± SD (Representative experiment with *n* = 3 technical replicates). **e.** Immunoblot analysis of isolated EVs and NVEPs. **f**. Nanoflow cytometry analysis of CD63 marker on isolated EVs and NVEPs. **g**. Representative images of iTF-microglia under control, pro-inflammatory or anti-inflammatory conditions stained for mitochondria marker COXIV (red) and F-actin filaments with phalloidin (green). Nuclei were labelled by DAPI. Scale bar: 10 μm. **h**. Quantitative RT-PCR analysis of pro-inflammatory markers MCP-1 and IL-6 and anti-inflammatory markers Arginase-1, CD206 and IL-10 mRNA expression in differentiated iTF-microglia treated with LPS, IL-10 or control conditions. Rsp18 was used as an endogenous control to normalize gene expression levels. Relative mRNA abundance in control to LPS or IL-10 iTF-microglia is reported. Results represent mean ± SD (Representative experiment with *n* = 3 technical replicates). **i.** Bioanalyzer image of RNA quality extracted from sEV, exomeres and supermeres samples. **j.** Quantitative RT-PCR analysis of miR-146a-5p expression in EVs and NVEPs isolated from iTF-microglia treated with LPS, IL-10 or control conditions. Cel-miR-39 spike in was used as control to normalize miRNA expression levels. Relative miRNA abundance in control to LPS or IL-10 iTF-microglia EVs and NEVPs is reported. Results represent mean ± SD (Representative experiment with *n* = 3 technical replicates).

### LPS-induced neuroinflammation in mice alters microglial activation, EV and NVEP profiles, and miR-146 expression

To provide *in vivo* validation of our previous findings *in vitro*, mice were administered LPS (0.5mg/kg) injected IP to induce neuroinflammatory conditions as previously described ^32,33^. To assess behavioral changes associated with neuroinflammation, we performed a marble burying test^34^. This assay is widely used as a reproducible measure of repetitive and anxiety-like behaviors, which are commonly influenced by neuroinflammatory processes. Because neuroinflammation can disrupt neural circuits involved in motivation, stress response, and compulsive behavior, changes in marble burying activity serve as an indirect yet reliable indicator of neuroinflammatory states *in vivo*^35^. LPS-treated animals exhibited a reduction in marble burying behavior, reflecting decreased exploratory drive and motivation, consistent with sickness behavior and anxiety-like states associated with neuroinflammation. (Fig. 5a). qRT-PCR analysis revealed increased expression (in mouse brain) of microglial activation markers (Clec4d, Msr1, Marco) and decreased expression of homeostatic microglial markers (Fcrls, P2ry12) (Fig. 5b) consistent with a transition toward a pro-inflammatory microglial state. In parallel, the upregulation of inflammatory cytokines (Il1b, TNF, Nos2) (Fig. 5c) and the astrocytic activation marker Gfap (Fig. 5d) indicates coordinated activation of glial populations. Together, these cell-type–specific transcriptional changes provide strong evidence of a neuroinflammatory response.

**Figure 5:**
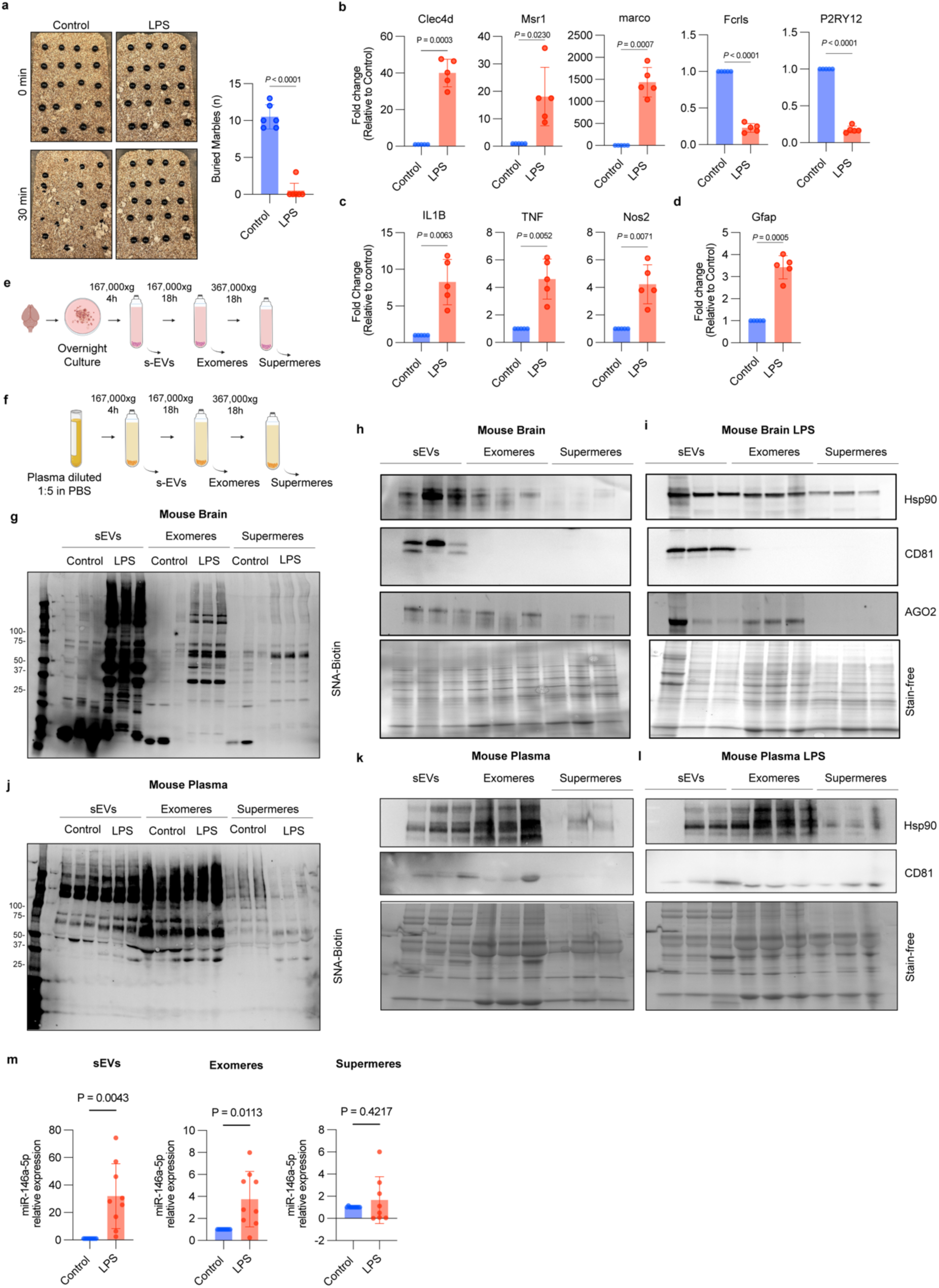
Characterization of EVs and NVEPs in an LPS-Induced Neuroinflammation Mouse Model. **a.** Representative images of the experimental cages before and after marble burying test. Numbers of marbles buried quantification (*n* = 6 mice per group). **b**. Quantitative RT-PCR analysis of associated microglia activation markers (Clec4d, Msr1, Marco) and homeostatic microglia markers (Fcrls, P2ry12). **c**. Quantitative RT-PCR analysis of inflammatory markers (Il1b, TNF, Nos2). **d**. Quantitative RT-PCR analysis of the astrocytic marker Gfap. Rsp18 was used as an endogenous control to normalize gene expression levels. Relative mRNA abundance is reported. Results represent mean ± SD (Representative experiment with *n*=5 mice per group and *n* = 3 technical replicates). **e**. Schematic representation of the mouse brain-derived EVs and NVEPs isolation protocol. **f**. Schematic representation of the mouse plasma-derived EVs and NVEPs isolation protocol. **g.** Representative images of *Sambucus nigra agglutinin* (SNA) blot analysis of mouse brain EVs (large and small EVs) and NVEPs (exomeres and supermeres). **h.** Immunoblot analysis of isolated brain EVs (large and small EVs) and NVEPs (exomeres and supermeres). **i.** Immunoblot analysis of isolated LPS-brain EVs (large and small EVs) and NVEPs (exomeres and supermeres).**j.** Representative images of SNA blot analysis of mouse plasma EVs (large and small EVs) and NVEPs (exomeres and supermeres). **k.** Immunoblot analysis of isolated plasma EVs (large and small EVs) and NVEPs (exomeres and supermeres). **l.** Immunoblot analysis of isolated LPS-plasma EVs (large and small EVs) and NVEPs (exomeres and supermeres).**m**. Quantitative RT-PCR analysis of miR-146a-5p expression in EVs and NVEPs isolated from plasma-derive EVs and NVEPs under LPS or control conditions. Cel-miR-39 spike in was used as control to normalize miRNA expression levels. Relative miRNA abundance in control to LPS EVs and NEVPs is reported. Results represent mean ± SD (Representative experiment with *n* = 8 mouse per group and *n* = 3 technical replicates).

Prior investigators have successfully isolated EVs from brain tissue^36,37^. We adapted this methodology to isolate EVs and NVEPs from mouse brain tissue by ultracentrifugation^38^ (Fig. 5e). As previously noted in our *in vitro* models (Fig 2h), SNA lectin staining revealed distinct α2,6-sialylation patterns, with increased signal detected in brain sEVs, exomeres and supermeres upon LPS treatment (Fig. 5g). Immunoblot analysis confirmed enrichment of canonical EV marker CD81 in sEV fractions, whereas exomeres and supermeres exhibited distinct protein profiles (Fig. 5h-i). EVs, exomeres and supermeres were isolated from plasma (Fig. 5f), characterized by immunoblotting (Fig. 5k-l) and stained with lectin. Protein concentration was measured; exomeres had the highest protein concentration and no differences were observed between male and female mice (Extended Fig. 5a and 5b). Unlike brain-derived particles, SNA lectin staining for plasma-derived EVs and NVEPs did not show distinct α2,6-sialylation patterns for the different states (Fig. 5j). To determine whether plasma-derived sEVs or NVEPs carry our candidate microRNA miR-146a-5p linked to microglial inflammatory state, we quantified its levels by qRT-PCR. LPS stimulation led to a marked enrichment of miR-146 in both sEVs and exomeres compared to controls, with the strongest signal detected in small EVs (Fig. 5m). Furthermore, by utilizing the CNS microRNA profiles data base^39,40^, which compiles comprehensive expression data of microRNAs in the central nervous system, we identified a significant upregulation of mmu-miR-146a-5p in microglia located within the brainstem (Extended Fig. 5c).

### Microglia-derived EVs are found in circulation and transport inflammatory miRNA-146a-5p cargo

To directly capture microglia-derived EVs (rather than bulk EVs) in the plasma in vivo, we utilized our *Exomap1* transgenic mouse model^41^, which encodes hCD81-mNeonGreen fusion protein integrated into the *H11* locus downstream of a lox-stop-lox (LSL) cassette^41^ (Fig.6a). By crossing the *Exomap1* mice with *Fcrls-2A-Cre* mice^42^, which expresses Cre recombinase under microglia cell-type-specific promotor Fcrls, we generated mice that express the hCD81-mNeonGreen selectively in microglia cells and in the EVs derived from those cells. Flow cytometry of isolated mouse primary microglia confirmed the robust expression of mNeonGreen (mNG) upon Cre-mediated recombination (Fig. 6b). We next assessed whether microglia-derived EVs could be detected in circulation. EVs isolated from plasma of *Fcrls-2A-Cre×Exomap1* mice exhibited mNG signal as detected by nanoflow cytometry, whereas EVs from control *Exomap1* mice lacked detectable fluorescence (Fig. 6c e). Further, SEVEN reveled mNG signal in hCD81-enriched EVs isolated from plasma of *Fcrls-2A-Cre×Exomap1* mice (Fig. 6d, left). To confirm the brain origin of these vesicles, we used SEVEN to analyze EVs-derived from brain tissue in *Fcrls-2A-Cre×Exomap1* mice (Fig. 6d, right). We observed a significant number of hCD81-mNG containing EVs in these mice (n=3), while negligible signal was detected from control *exomap1 mice* (n=3) (Fig. 6e, Table S4). Characterization of biophysical and molecular features for hCD81-mNG-enriched EVs revealed a heterogenous size distribution, with an average size of 101 nm (Fig. 6f); and enrichment of TSPAN cargo molecules, with an average of 20 detected TSPAN molecules per vesicle (Fig. 6g). These EVs were predominantly circular, with an average circularity of 0.84 (Fig. 6h). Descriptive statistics for EV properties are shown in Table S5.

**Figure 6:**
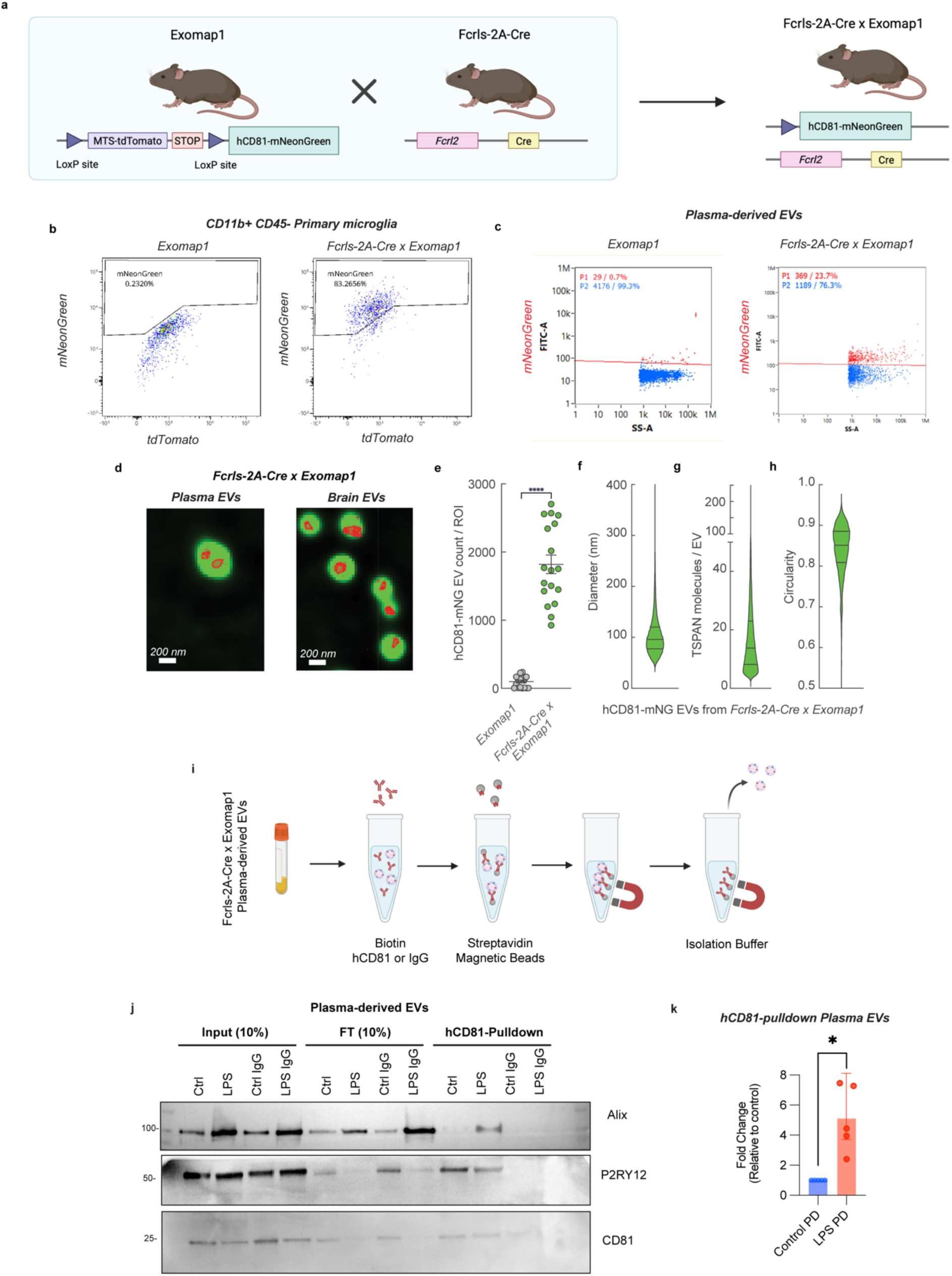
Characterization of microglia-derived EVs from cell-type specific hCD81-mNeonGreen mouse model. **a**. Schematic representation of microglia cell-type-specific engineered extracellular vesicles mouse model. **b**. Flow cytometry staining of CD11b+ CD45- primary microglia to assess the colocalization of mNeonGreen (hCD81-mNeonGreen) (*n*=3 mice per group). **c**. Nanoflow cytometry of plasma-derived EVs. **d.** Images of hsCD81-mNG EVs from plasma and brain tissue. SMLM signal detecting tetraspanins is in red (used for quantification) and SRRF signal (detecting mNeonGreen) is in green. **e**. Average number of detected hsCD81-mNG EVs per ROI detected from tissue of Exomap1 and *Fcrls-2A-CrexExoma1* mice (*n*=3 mice per group, 5 ROIs per mice). Mean ± SEM; ****p<0.0001. **f**. Size distribution for all detected hcCD81-mNG EVs from *Fcrls-2A-CrexExoma1* mice (*n*=3 mice). **g** TSPAN number per EV for all detected hcCD81-mNG EVs from Exomap1 Fcrls-2A-Cre mice (*n*=3 mice). **h.** Circularity for all detected hcCD81-mNG EVs from Exomap1 Fcrls-2A-Cre mice (*n*=3 mice). Median thick horizontal line, quartiles thin horizontal line, crosses averages. Descriptive statistics and significances are shown in Tables S4 and S5. **i**. Schematic representation of experimental pipeline for isolating microglia cell-type-specific extracellular vesicles from mouse plasma. **j**. Immunoblot analysis of extracellular vesicles from microglia, Alix and CD81 (EV markers) and P2RY12 (microglia marker). **k**. Bar plot showing significant differences in miR-146a-5p between humanized CD81-positive EV pulldown in control vs LPS treated mouse, with respective p-values using Mann Whitney t-test (*n*= 5 mice per group). Mean ± SEM; ****p<0.0001.

To selectively isolate microglia-derived EVs from plasma, we developed an immunocapture strategy targeting hCD81 expressed in the EVs released from Cre-expressing microglia (Fig. 6i). Immunoblotting of hCD81-captured EVs confirmed the presence of the microglial marker P2RY12 and canonical EV markers Alix and CD81 in both control and the neuroinflammation model with LPS treatment (Fig. 6j), although the level of Alix was higher in the LPS-treated group. Finally, we verified that miR-146a-5p, that we identified as an EV biomarker for microglial inflammation, was markedly increased in the plasma microglial-derived EVs in neuroinflammation compared to control mice (Fig. 6k). These in vivo data were strongly supportive of a novel EV-RNA based biomarker associated with microglial inflammation that could be detected in the plasma.

### Immunoaffinity capture optimally facilitates specific characterization of EVs and NVEPs from human plasma

To translate our previous findings into a clinically relevant context, we next evaluated the feasibility of isolating and profiling these distinct particles in human plasma. Human plasma-derived EVs and NVEPs were isolated according to the procedures outlined in Fig. 7a. To characterize the size distribution, isolated EVs (large and small) and NVEPs (exomeres and supermeres) were imaged by transmission electron microscopy, which revealed distinct size differences, with NVEPs smaller than sEVs (Fig.7b). Using SEVEN^35^ we further characterized EVs and NVEPs (Fig. 7c, g and k) from crude plasma and isolated plasma fractions (sEVs, exomeres, and supermeres). As with HMC3 cells, we used TSPAN capture/detection for EVs and SNA capture/detection for NVEPs and additionally evaluated SNA-enriched EVs (SNA capture/TSPAN detection). Crude plasma and sEV fractions contained many TSPAN-enriched EVs, whereas exomere and supermere fractions yielded markedly fewer particles (Fig. 7d, Table S6) suggesting that the ultrafiltration/differential centrifugation isolation method successfully depletes EVs from the exomere and supermere fraction. The presence of TSPAN-positive particles in exomere and supermere fractions may be consistent either with the presence of TSPAN on these particles, or the co-isolation of some sEVs. Although more abundant in crude plasma, SNA-enriched EVs were rare in all fractions (Fig. 7h, Table S6). Compared to EVs, SNA-captured NVEPs were significantly more abundant across all fractions, with crude plasma containing ∼4–10-fold more NVEPs than the isolated fractions (Fig. 7l, Table S6). Of note, while exomere and supermere fractions show substantial EV depletion compared to sEV fraction, SNA-enriched NVEPs abundant in all three fractions, suggesting a significant degree of co-isolation of NVEPs with the sEV fraction (potentially leading to confounding effects in biomarker studies).

**Figure 7:**
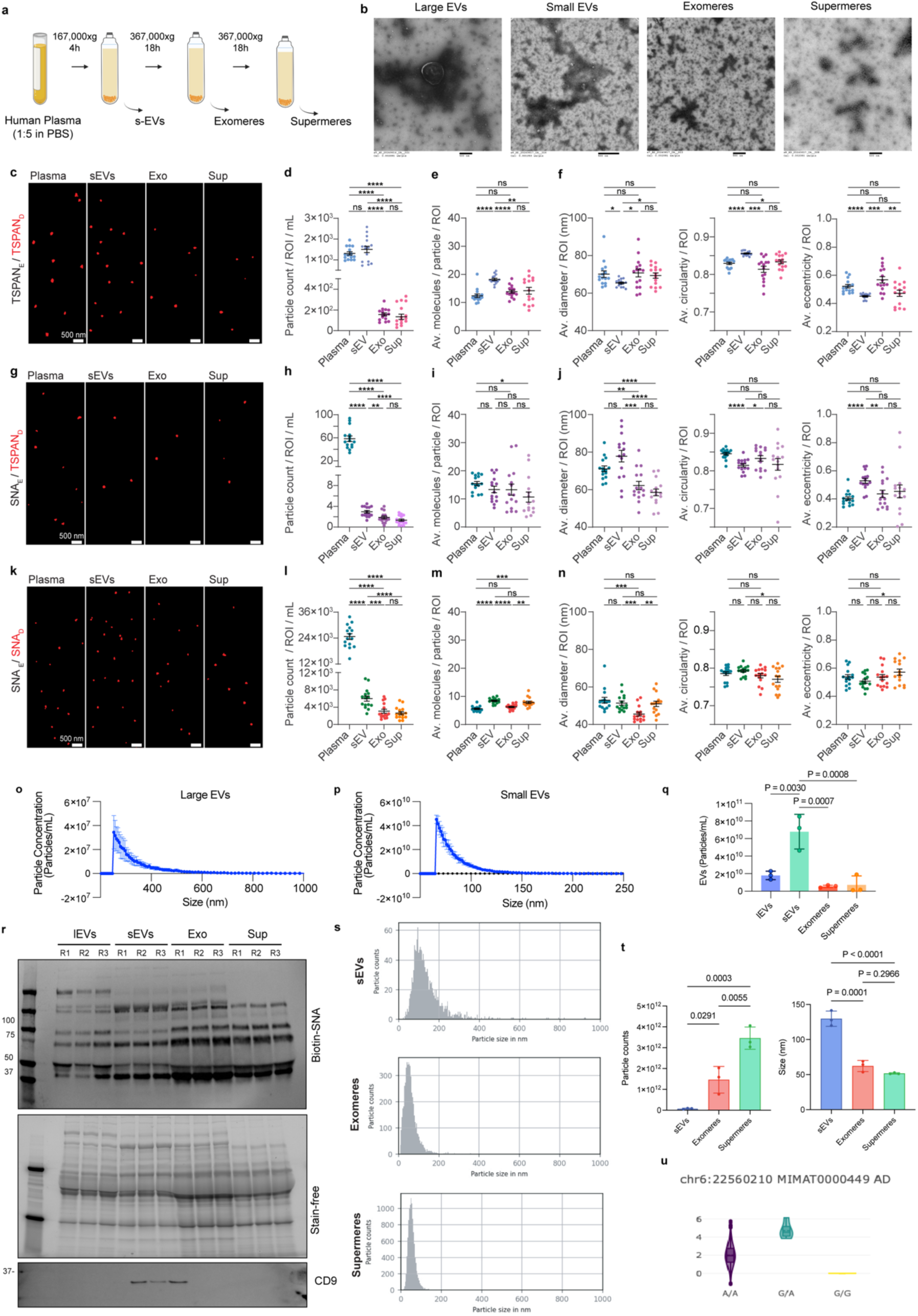
Extracellular vesicles and non-vesicular extracellular nanoparticles can be isolated from human plasma and used as predictors of Alzheimer’s disease. **a.** Schematic representation of the sEV and NVEPs (exomeres and supermeres) isolation protocol. **b.** Transmission electron microscopy (TEM) analysis of human pool plasma-derived large EVs, small EVs, exomeres and supermeres. **c.g.k.** Representative images of TSPAN-enriched EVs (c), SNA-enriched EVs (g), and SNA-enriched NVEPs (k) from crude plasma and isolated EV, exomere and supermere fractions (dilutions were not uniform and adjusted to optimize particle sampling). Scale bar 500 nm. **d.h.l.** Number of detected particles for crude plasma and isolated fractions (sEVs, exomere, and supermere). Data was normalized per 1 μl of plasma.**e.i.m.** Number of detected cargo molecules per particle (TSPANs for EVs and SNA detected glycans for NVEPs) for crude plasma and isolated fractions (sEVs, exomere, and supermere). **f**.**j.n.** Quantification of morphology: size (left), circularity (middle), eccentricity (right) for crude plasma and isolated fractions (sEVs, exomere, and supermere). Averages ± SEM are shown (n=3 independent experiments, each imaged with 5 ROI). **o.** Microfluidic resistive pulse sensing (MRPS) data of isolated large EVs. Data represented as Mean ± SD (n=3 samples per group). **p**. Microfluidic resistive pulse sensing (MRPS) data of isolated small EVs. Data represented as Mean ± SD (n=3 samples per group). **q**. Particle concentration in the isolated EVs and NVEPs quantified by TSPAN ELISA (N=3 samples per group) (Data are mean particles/ml ± SD). **r**. Representative images of SNA blot analysis of cell lysates, EVs (large and small EVs) and NVEPs (exomeres and supermeres). **s**. Size distribution of plasma-derived EVs, exomeres and supermeres in scatter mode using NTA. **t**. Particle concentration and median Size measured by NTA. Data represented as Mean ± SD (n=3 samples per group). **u**. Association plot of has-miR-146a-5p (chromosome 6, MAF 22560210, A/G, p-value 1.23e-7) with Alzheimer’s disease. Japanese miRNA-eQTL database.

We next evaluated particle cargo and morphology of the detected particles. Relative to crude plasma, sEVs from isolated fraction displayed higher TSPAN molecules content and appeared slightly smaller, more circular, and less elongated (Fig. 7d,f); this is consistent with corona molecules (EV-associated plasma factors)^10, 11^ on EVs from crude plasma that were largely removed with the isolation protocol. Isolated exomere and supermere fractions characterized by TSPAN capture/detection showed similar properties to those found in crude plasma (Fig. 7e, f). SNA-enriched EVs had similar TSPAN content across different fractions (Fig. 7i), but their size was slightly but significantly higher in crude plasma and sEV fraction compared to exomere and supermere fractions (Fig. 7j). SNA capture/detection of NVEPs from crude plasma (compared to sEV isolated fraction) resulted in fewer SNA-detected glycans (Fig. 7m); both had average sizes slightly above 50 nm, average circularities under 0.8, and average eccentricities above 0.5 (Fig. 7n). Notably, SNA capture/detection of NVEPs from supermeres and exomere fractions showed some differences. NVEPs from exomere fractions (compared to supermere fractions) displayed fewer detected glycans (6 vs. 8) and slightly smaller diameters (46 nm vs. 51 nm) compared to supermeres, likely reflecting the greater aggregation propensity of plasma supermeres (Fig. 7m). As expected for non-vesicular entities, NVEPs exhibited low circularity and high eccentricity (Fig. 7n). Descriptive characteristics for properties are listed in Table S7 (averages per ROI) and Table S8 (all individual EPs).

Crude plasma contains abundant lipoprotein particles (LPPs), with ApoE highly represented across several LPP subclasses with sizes comparable to EVs/NVEPs (Extended Fig.7a). Additionally, we check ApoA levels in isolated sEVs, exomeres and supermeres (Extended Fig.7b), as is often present on the EV corona^43^. We further characterized ApoE-enriched EVs and LPPs from crude plasma as APOE is also found in EV corona (Extended Fig. 7d-e)^43^. As expected, ApoE-enriched LPPs were significantly more abundant compared to ApoE-enriched EVs (Extended Fig. 7f, Table S9). ApoE-enriched EVs contained an average of 18 detected TSPAN molecules (Extended Fig. 7g), had average size of 61 nm (Extended Fig. 7h), and were largely circular, consistent with canonical EVs (Extended Fig. 7i). ApoE-enriched LPPs had sizes similar to exomeres and supermeres (∼49 nm on average), but they had a different shape compared to NVEPs; their average circularities were above 0.8 and eccentricities below 0.5 (Extended Fig. 7j). Descriptive characteristics for properties are listed in Table S10. Altogether, by capturing/detecting different molecules and examining their cargo content, size, and shape, we can clearly distinguish between main particle types present in crude plasma.

We next characterized ^44,45^large and small sEV size using another technique microfluidic resistive pulse sensing (MRSP), based on changes in electrical resistance as individual particles pass through a small fluidic constriction. MRSP revealed large EVs measured around 200-400 nm (Fig. 7o), while small EVs measured around 50-100 nm (Fig. 7p). Particle concentration in the isolated EVs and NVEPs was quantified by TSPAN ELISA, sEVs were enriched in tetraspanins (Fig. 7q). Lectin staining revealed distinct α2,6-sialylation patterns in plasma EVs and NVEPs (Fig. 7r). EVs and NVEPs were further characterized by immunoblotting (Extended Fig. 7c). We detected the presence of canonical EV markers (membrane EV markers: CD81 and NVEP-associated markers (ACE2 and LDHA). Additionally, we used nanoparticle tracking analysis to measure particle concentration and size in isolated sEVs, exomeres and supermeres (Fig. 7s). We found that supermeres had higher concentration in comparison to exomeres and sEVs (sEVs: 7.8×10^10^ ± 2.49×10^10^ particles/mL; exomeres: 1.46×10^12^ ± 6.41×10^11^ particles/mL; supermeres: 3.46×10^12^ ± 5.35×10^11^ particles/mL) (Fig. 7t). In contrast, sEVs presented larger sizes in comparison to exomeres and supermeres (sEVs: 129.7 ± 10.88 nm; exomeres: 62.23 ± 7.905 nm; supermeres: 51.73 ± 0.9238 nm) (Fig. 7t). In totality, our detailed characterization of human plasma suggested that while the currently utilized isolation protocol depletes EVs from the exomere and supermere fractions, significant amounts of NVEPs may remain in the sEV fraction. Immunoaffinity methods (such as TSPAN capture) prior to cargo analysis may improve the specificity of measuring sEV-specific cargoes.

Previous studies support the utility of measuring the cargo of EVs and NVEPs in plasma as putative biomarkers. Table 5 summarizes previous clinically relevant findings in the literature that shows dysregulation of hsa-miR-146a-5p across different cohorts and diseases (measured in bulk plasma). Notably, the variability in findings suggest that carrier-specific analysis of molecular cargo, and attention to the disease stage using the tools we have described may be needed to improve the rigor and reproducibility of these studies. Interestingly, the Japanese miRNA-eQTL database (https://jamir-eqtl.org/) several eQTLs of hsa-miR-146a-5p have been associated with Alzheimer’s disease (Fig. 7u), providing genetic validation for further characterization of this marker.

**Table 5:**
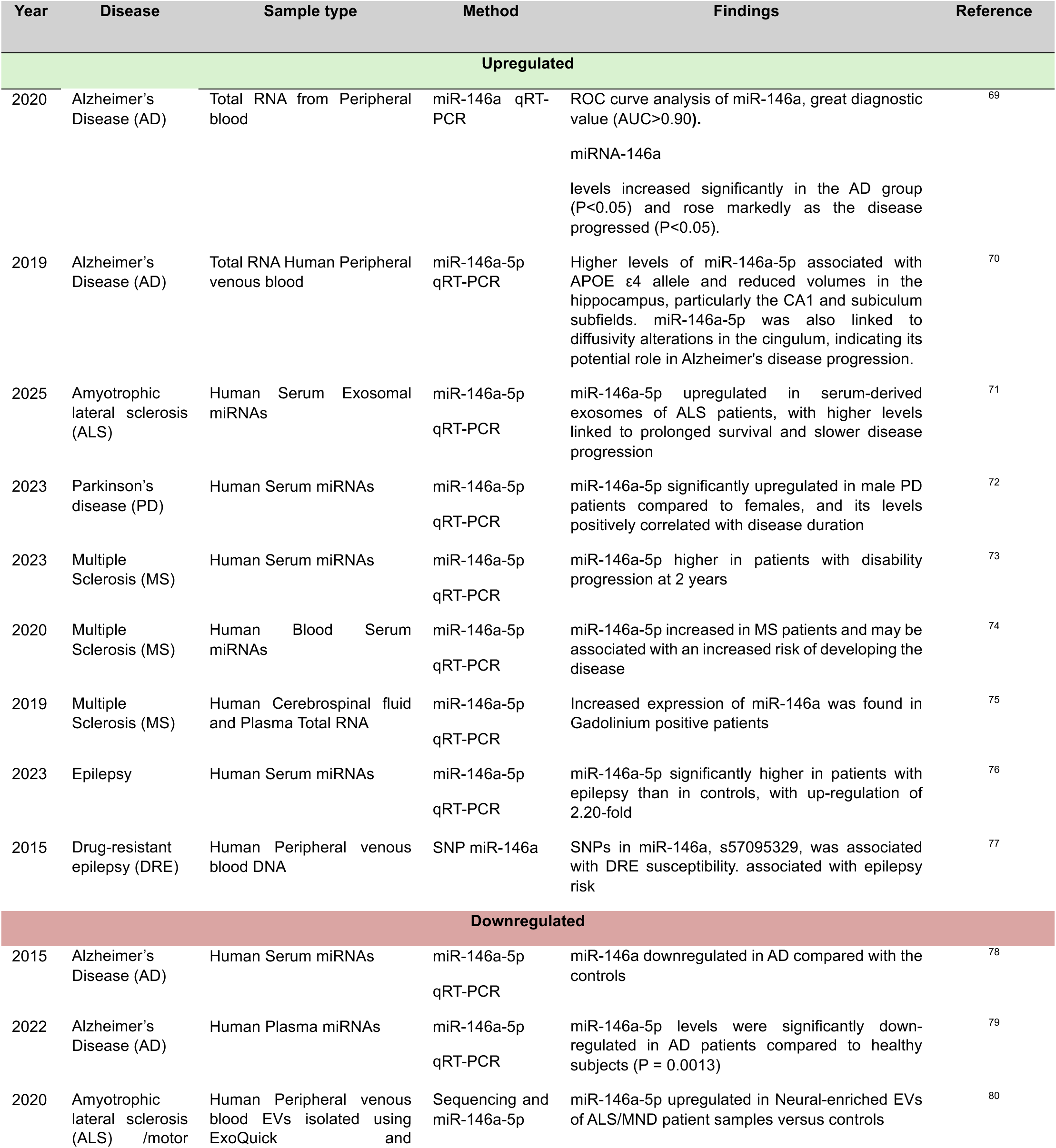
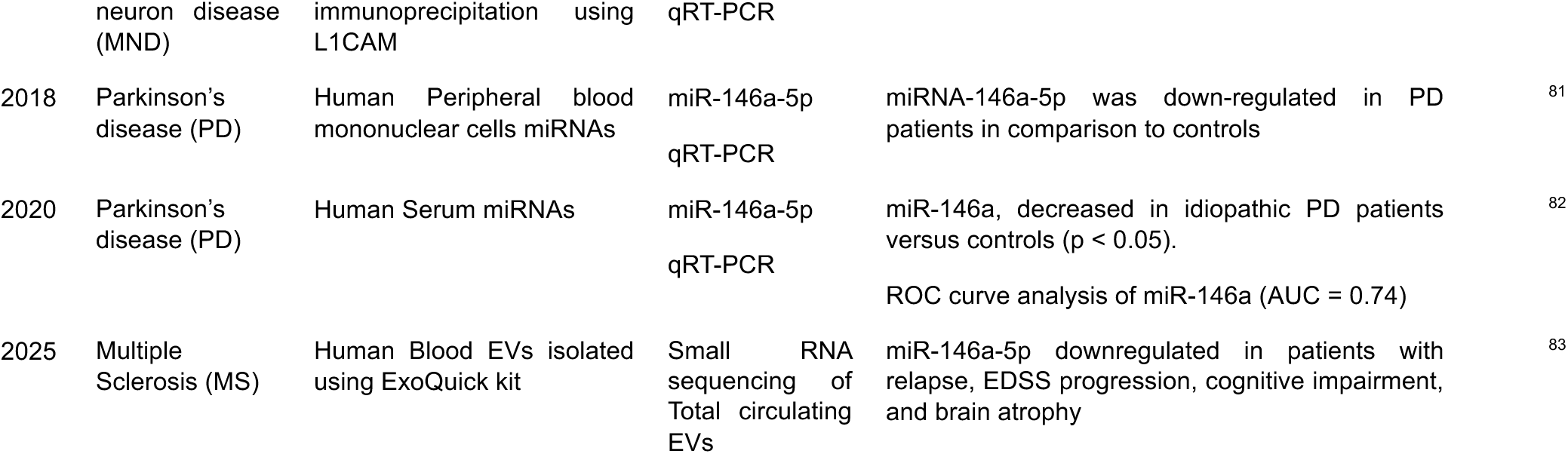
Summary of findings on miR-146a-5p as biomarker of human neurological disorders.

| Year | Disease | Sample type | Method | Findings | Reference |
| --- | --- | --- | --- | --- | --- |
| Upregulated |  |  |  |  |  |
| 2020 | Alzheimer's Disease (AD) | Total RNA from Peripheral blood | miR-146a qRT-PCR | ROC curve analysis of miR-146a, great diagnostic value (AUC>0.90).<br>miRNA-146a levels increased significantly in the AD group (P<0.05) and rose markedly as the disease progressed (P<0.05). | 69 |
| 2019 | Alzheimer's Disease (AD) | Total RNA Human Peripheral venous blood | miR-146a-5p qRT-PCR | Higher levels of miR-146a-5p associated with APOE ε4 allele and reduced volumes in the hippocampus, particularly the CA1 and subiculum subfields. miR-146a-5p was also linked to diffusivity alterations in the cingulum, indicating its potential role in Alzheimer's disease progression. | 70 |
| 2025 | Amyotrophic lateral sclerosis (ALS) | Human Serum Exosomal miRNAs | miR-146a-5p qRT-PCR | miR-146a-5p upregulated in serum-derived exosomes of ALS patients, with higher levels linked to prolonged survival and slower disease progression | 71 |
| 2023 | Parkinson's disease (PD) | Human Serum miRNAs | miR-146a-5p qRT-PCR | miR-146a-5p significantly upregulated in male PD patients compared to females, and its levels positively correlated with disease duration | 72 |
| 2023 | Multiple Sclerosis (MS) | Human Serum miRNAs | miR-146a-5p qRT-PCR | miR-146a-5p higher in patients with disability progression at 2 years | 73 |
| 2020 | Multiple Sclerosis (MS) | Human Blood Serum miRNAs | miR-146a-5p qRT-PCR | miR-146a-5p increased in MS patients and may be associated with an increased risk of developing the disease | 74 |
| 2019 | Multiple Sclerosis (MS) | Human Cerebrospinal fluid and Plasma Total RNA | miR-146a-5p qRT-PCR | Increased expression of miR-146a was found in Gadolinium positive patients | 75 |
| 2023 | Epilepsy | Human Serum miRNAs | miR-146a-5p qRT-PCR | miR-146a-5p significantly higher in patients with epilepsy than in controls, with up-regulation of 2.20-fold | 76 |
| 2015 | Drug-resistant epilepsy (DRE) | Human Peripheral venous blood DNA | SNP miR-146a | SNPs in miR-146a, s57095329, was associated with DRE susceptibility. associated with epilepsy risk | 77 |
| Downregulated |  |  |  |  |  |
| 2015 | Alzheimer's Disease (AD) | Human Serum miRNAs | miR-146a-5p qRT-PCR | miR-146a downregulated in AD compared with the controls | 78 |
| 2022 | Alzheimer's Disease (AD) | Human Plasma miRNAs | miR-146a-5p qRT-PCR | miR-146a-5p levels were significantly down-regulated in AD patients compared to healthy subjects (P = 0.0013) | 79 |
| 2020 | Amyotrophic lateral sclerosis (ALS) /motor | Human Peripheral venous blood EVs isolated using ExoQuick and | Sequencing and miR-146a-5p | miR-146a-5p upregulated in Neural-enriched EVs of ALS/MND patient samples versus controls | 80 |
| 2018 | Parkinson's disease (PD) | Human Peripheral blood mononuclear cells miRNAs | miR-146a-5p qRT-PCR | miRNA-146a-5p was down-regulated in PD patients in comparison to controls | 81 |
| 2020 | Parkinson's disease (PD) | Human Serum miRNAs | miR-146a-5p qRT-PCR | miR-146a, decreased in idiopathic PD patients versus controls ($p < 0.05$).<br>ROC curve analysis of miR-146a (AUC = 0.74) | 82 |
| 2025 | Multiple Sclerosis (MS) | Human Blood EVs isolated using ExoQuick kit | Small RNA sequencing of Total circulating EVs | miR-146a-5p downregulated in patients with relapse, EDSS progression, cognitive impairment, and brain atrophy | 83 |

## Discussion

Neuroinflammation is a hallmark of numerous neurological diseases and injuries, yet detecting and monitoring it within the brain remains a major clinical challenge. Reliable biomarkers of neuroinflammation are urgently needed to enable early detection, patient stratification, and monitoring of disease progression across conditions such as Alzheimer’s disease, multiple sclerosis, traumatic brain injury, and neuropsychiatric disorders^46^. Current diagnostic tools largely capture structural or metabolic changes that occur after irreversible neuronal loss, underscoring the need for molecular biomarkers that reflect earlier events^47^. Microglia, as the brain’s resident immune cells, are central regulators of neuroinflammatory responses^2,48^. Their activation precedes neuronal dysfunction and contributes to both disease onset and progression of neurodegenerative diseases^49^.Therefore, identifying microglia-derived molecular signals particularly those released into circulation offers a powerful approach to detect and monitor neuroinflammation noninvasively. In this study, we examined microglial activation under defined pro- and anti-inflammatory conditions and identified a microRNA that is robustly associated with the inflammatory state. Such a marker holds promise not only for improving our understanding of microglial dynamics, but also for enabling the development of minimally invasive diagnostics, particularly if it can be detected in accessible biofluids like cerebrospinal fluid or blood. A tool of this kind could provide critical insight into disease mechanisms, treatment response, and patient-specific trajectories in a broad range of CNS disorders.

EVs and newly discovered NVEPs are mediators of interorgan crosstalk that have been implicated in the pathogenesis of different diseases^12,21,25^. In this study, we demonstrated that microglia release distinct EVs and NVEPs under different activation states. Transmission electron microscopy, atomic force microscopy, and super-resolution microscopy revealed nanometer-scale differences, supporting the notion that NVEPs represent distinct entities rather than vesicular contaminants in EV isolations consistent with prior reports in cancer cells^22,24,25^. Immunoblotting characterization further showed that EVs and NVEPs carry unique molecular markers. The sialylation of glycoproteins in immune cells is critical for regulating immune responses^50,51^. N-linked glycosylation has distinct patterns in the mouse brain, with enrichment in choroid plexus harbors which harbors a variety of immune cells including microglia^52^. Lectin staining revealed different glycan α2,6-sialylation patterns which differentiate the NVEPs from canonical EVs. Moreover, we observed different glycan α2,6-sialylation patterns between isolated fractions and crude plasma, consistent with previous findings using glycan arrays^53^, suggesting that isolation methods might affect glycan measurements. These is not surprising since recently it has been shown that isolation methodology can modify NVEP composition^24^.

Recent studies have also found glycoRNAs (RNAs modified with secretory N-glycans) in the surface of different cell types including immune cells^54–56^, in EVs^57^ and NVEPs^58^. GlycoRNA profiles change during immune cell differentiation and activation, implicating their potential roles in innate immune responses^56^. However, despite these intriguing associations, the functional consequences of RNA glycosylation remain unknown. To date, no direct mechanistic role has been demonstrated for glycoRNAs in intercellular signaling or immune modulation. Based on this, we hypothesize that glycoRNAs may contribute to the observed differences in EVs and NVEPs. Notably, previous studies have reported an increase in α2,6-sialylation in aging and AD, particularly in plaque-associated microglia^59^. This aligns with our findings in microglia EVs and NVEPs under pro-inflammatory activation, suggesting that altered α2,6-sialylation may represent a shared mechanism linking microglial activation and neuroinflammatory signaling. Single particle analysis of EVs and NVEPs using SEVEN enabled us to obtain their multiparametric properties. For the first time, we were able to obtain detailed morphology (size with shape) for supermeres and exomeres. As expected, NVEPs (compared to EVs) were smaller and had less circular shape due to their non-vesicular nature. The data also revealed unique signatures for pro-inflammatory vs anti-inflammatory treatments of EPs. Pro-inflammatory and anti-inflammatory treatments significantly affected EVs, modestly affected exomeres, and marginally affected supermeres. Additionally, the effect of anti-inflammatory treatment (compared to pro-inflammatory) had more profound effect on EP concentration and morphology. Our small RNA-sequencing data revealed that EVs and NVEPs released during different microglia activation states contain differentially expressed small RNAs. Although previous studies have reported circulating RNAs associated with neurodegenerative and inflammatory processes^60,61^, to our knowledge, this is the first systematic and detailed assessment of the RNA cargo of EVs and NVEPs derived from microglial cells. Our data revealed miRNAs as the most diverse class of small RNA cargoes in EVs and NVEPs. Among all the miRNAs, miR-146a-5p emerged as a consistent and robust marker of pro-inflammatory activation across species and experimental models in our study. We observed its upregulation in EVs and to a lesser degree in exomeres released from pro-inflammatory microglia in HMC3 cells, human iPSC-microglia and mouse plasma-derived EVs. To prove the microglial origin of plasma miR-146 in our model of neuroinflammation, we leveraged our Fcrl-2a-cre- *Exomap1* double transgenic system, which corroborated the overexpression of miR-146a-5p in microglia-derived EVs isolated from plasma upon LPS-induced neuroinflammation. Together our cellular and murine data confirmed the presence of distinct biophysical and RNA cargo differences in EVs and exomeres and converged on EV-miRNA 146 as a promising peripheral marker for neuroinflammation.

To translate our findings toward clinical relevance, we next examined whether these molecular entities could be characterized in isolation (without confounding co-isolation) in human plasma, a biological context where they remain largely unexplored. We extended our characterization of EVs, exomeres, and supermeres by analyzing both isolated fractions and plasma samples directly, providing a more clinically pertinent assessment. Remarkably, we discovered that supermeres represent the most abundant particle population in human plasma even compared to ApoE-enriched lipoproteins, suggesting a potentially untapped source of biomarkers. Alternatively, given their relative abundance, and the persistent presence of NVEPs in sEV fractions, they may act as potential confounders in biomarker studies. Reassuringly, our immunoaffinity-based imaging methodologies (e.g. using TSPAN or SNA capture) are able to efficiently study the characteristics and cargo of these particles in isolation. Since sEVs contribute to a small portion of the particles in plasma, immunoaffinity-based technologies to isolate microglia-specific EVs in plasma may markedly enhance the signal for monitoring neuroinflammatory changes non-invasively. Here, we have already shown that hCD81-enrichment of EVs from Fcrls-2A-CrexExomap1 mouse model (microglial origin) allowed the detection of increase levels of miR-146a-5p upon pro-inflammatory treatment. Further, building on our previous work identifying EV cargo using the SEVEN platform^62^, we anticipate that this approach will also enable the detection and characterization of microRNAs within these particles.

Circulating EVs in human plasma from patients with neurodegenerative diseases, have been shown to have dysregulated levels of miR-146a-5p (Table 5). However, these studies, conducted mostly in total plasma have yielded inconsistent results. While some of the variability may reflect the stage of disease and potential peripheral feedback loops, the lack of measurement of miR-146a-5p in sEVs or more specifically, in microglial-derived sEVs likely is a more significant contributor to the disparate results. Our current study robustly demonstrates the presence of this miRNA in EVs and exomeres released by microglial cells in the acute stage of inflammation.

Together, these findings strongly implicate microglia-derived extracellular particles as dynamic, state-dependent markers of neuroinflammation that can be detected in the periphery. The detection of microglial EVs in the plasma and measurement of their RNA cargoes show promise as potential non-invasive biomarkers to complement emerging biomarkers such as amyloid-beta and tau protein in diagnosing pre-symptomatic AD as well as other diseases in which neuroinflammation plays a role.

### Limitations and future directions

Despite the promising findings, several limitations should be acknowledged in this study. One limitation is the use of ultracentrifugation methods (UC) for the isolation of EVs and NVEPs. UC is labor-intense and time consuming, requiring several days using specialize equipment. Furthermore, the differential centrifuge forces used, co-isolate contaminants that are pelleted on the same speed, which makes challenging to obtain pure samples. These high centrifuge forces also cause particle aggregation which might influence their molecular properties and strip the corona from EVs. Gentler and more specific methods (such as immunoaffinity isolation of these particles) may overcome these limitations.

Another limitation is that most of our findings are *in vitro* and murine models of microglial activation, which may not fully capture the complexity of human neuroinflammatory processes in Alzheimer’s disease (AD) or other diseases. Further validation in human-derived microglia from patients and larger, clinically diverse patient cohorts will be necessary to determine the translational relevance of the identified EV and NVEP markers. In addition, we demonstrated that miR-146a-5p is a consistent marker of pro-inflammatory activation, however its levels across different diseases and stages in relation to EVs and NVEPs remain to be fully elucidated. Moreover, the mechanisms by which altered α2,6-sialylation and glycoRNA profiles influence EV/NVEP biogenesis and cargo selection warrant deeper investigation, as these may represent novel regulatory layers in neuroimmune signaling.

Future studies integrating multi-omics and spatial mapping of microglial activation with circulating EV/NVEP signatures will further clarify their biological roles and clinical potential in relevant diseases context. Ultimately, new methods to isolate and characterize microglia-derived extracellular particles will be key to establishing their utility as early, noninvasive biomarkers complementing current biomarkers for the diagnosis of neurodegenerative diseases.

## Materials and methods

### Cell Culture

HMC3 Cells (CRL-3304, ATCC) were cultured in DMEM (Catalog number 11995073, Gibco) supplemented with 10% fetal bovine serum (Catalog number A5669701, Gibco) and 1% penicillin and streptomycin (Catalog number 15140122, Gibco, Thermo Fisher Scientific). Cells were maintained at 37°C, 5% CO2 and 95% humidity in a CO_2_ incubator.

For extracellular vesicles and non-vesicular extracellular particle collection, cells were cultured at 80% confluency, wash with PBS and cultured in serum free media conditions for 48 hours.

### Human iPSC and iPSC-derived microglia

Human iPSCs (male WTC11 background, PMID 24509632) were obtained from the Kampmann lab^31^ at the University of California, San Francisco (UCSF) and cultured in StemFlex Basal Medium (Catalog number A33493-01, Gibco) on BioCoat Cell Culture Treated Dishes (Catalog number 356413, Corning) coated with growth factor reduced, phenol red-free, LDEV-Free Matrigel Basement Membrane Matrix (Catalog number 356231, Corning) diluted 1:100 in Knockout DMEM (Catalog number 10829-018, Gibco). StemFlex was replaced every other day or every day. When 70%-80% confluent, cells were passaged by aspirating media, washing with D-PBS (Catalog number 14190-144, Gibco), incubating with StemPro Accutase Cell Dissociation Reagent (Catalog number A11105-01, Gibco) at 37°C for 5 min. After cells were collected in conical tubes and diluted 1:5 in StemFlex and centrifuged at 220xg for 5 min. Then cell pellets were resuspended in StemFlex supplemented with 10nM Y-27632 dihydrochloride ROCK inhibitor (Catalog number 125410, Tocris), cells were counted and plated onto matrigel-coated plates at the desired number.

Differentiation was performed following Kampmann lab’s protocol^31^. iPSCs were washed with D-PBS, dissociated with StemPro Accutase Cell Dissociation Reagent (Catalog number A11105-01, Gibco), and centrifuged at 220xg for 5 min. Cell pellets were resuspended in Essential 8 Basal Medium (Catalog number A1517001, Gibco) supplemented with 10 μM Y-27632 ROCK Inhibitor and 2 μg/mL doxycycline, then seeded onto double-coated plates with Poly-D-Lysine-precoated BioCoat plates and growth factor reduced, phenol red-free, LDEV-Free Matrigel Basement Membrane Matrix (Catalog number 356231, Corning). On day 2, cells were switched to differentiation medium (Advanced DMEM/F12 (Catalog number 12634010, Gibco) supplemented with 1× GlutaMAX (Catalo number 35050061, Gibco), 2 μg/mL doxycycline, 100 ng/mL human IL-34 (Catalog number 577904, BioLegend) and 10 ng/mL human GM-CSF (Catalog number 572904, BioLegend). On day 4, the medium was replaced with iTF-Microglia medium (Advanced DMEM/F12 supplemented with 1× GlutaMAX, 2 μg/mL doxycycline, 100 ng/mL human IL-34, 10 ng/mL human GM-CSF, 50 ng/mL human M-CSF (Catalog number 574804, BioLegend) and 50 ng/mL human TGF-β1 (Catalog number 781804, BioLegend). On day 8, fresh iTF-Microglia medium was added. Cells were maintained for 9 more days with full medium changes every 2–3 days.

### Lipopolysaccharide (LPS) and Interleukin-10 (IL-10) treatments

To model pro-inflammatory and anti-inflammatory conditions, HMC3 cells or iTF-microglia were treated for 48 hours with either 100 ng/mL of LPS (Catalog number 00-4976-03, Thermo Fisher Scientific) to induce a pro-inflammatory response, or 50 ng/mL of IL-10 (Catalog number 200-10, Prepotech) to induce an anti-inflammatory response.

### Mouse studies

All animal experiments were approved by the Institutional Animal Care and Use Committee at the Massachusetts General Hospital (2016N000047). Mice were maintained on a standard light–dark cycle with ad libitum access to food and water in a room with controlled temperature (22°C) and humidity (around 50%). Cages were cleaned every 4–5 days, and water and food supplies were checked daily. All mice were purchased from The Jackson Laboratory. We use C57BL/6J (Stock No: #000664) for LPS studies. To induce hCD81-mNeonGreen expression specifically in microglia cells, *exomap1* mice^41^ were crossed with mice expressing Cre recombinase driven by microglia specific promoter C57BL/6J-Fcrl2em1(cre)Gfng/J^42^ (Stock No: #036591). All strains were genotyped by Transnetyx, genotyping primers can be found on Table 2. For neuroinflammatory studies mice were injected intraperitoneally with Lipopolysaccharide from Escherichia coli (LPS) 0.5 mg/kg for 24 hours.

### Marbel burring test

Each mouse was placed in an individual cage with twenty marbles distributed in 5 rows on top of the bedding. Mice were allowed to explore the cage for 30 minutes; after the number of buried marbles was measured. Buried marbles were calculated as giving value 1 to buried and 0 to don’t buried marbles. Images were taken before and after 30 minutes.

### Human plasma samples

Human pooled plasma was commercially obtained (Catalog number CCN-10, Biologic) and reported to contain platelet-poor plasma from at least 20 healthy donors aged between 18 and 66.

### Extracellular vesicles and non-vesicular nanoparticles isolation from cell-conditioned media

Extracellular vesicles (EVs) and non-vesicular extracellular particles (NVEPs) were isolated from cell-conditioned medium as previously described with minor modifications^27^. Conditioned media from cell culture was collected and then centrifuged for 5 minutes at 500xg to remove cells. After it was centrifuged at 1,000xg for 20 minutes to remove debris. Then the supernatant was filtered through a 0.22 μm filter (Catalog number 229707, Celltreat) and concentrated using a 100,000 molecular-weight cutoff (Catalog number UFC710008, EMD Millipore). Concentrated media was the centrifuged at 10,000xg for 40 minutes and the pellet (lEVs) was resuspended in resuspended in PBS containing 25 mM HEPES (pH 7.2) (PBS-H). The supernatant was then centrifuged at 167,000xg for 4 hours in a SW32 Ti swinging-bucket rotor (Beckman Coulter) and the pellet (sEVs) was resuspended in resuspended in PBS-H. Then the supernatant was ultracentrifuged at 167,000g for 16 hours. The resulting pellet (exomeres) was resuspended in PBS-H. The supernatant was ultracentrifuged at 367,000g for 16 hours and the resulting pellet was resuspended in PBS-H (supermeres).

### Extracellular vesicles and non-vesicular nanoparticles isolation

Extracellular vesicles (EVs) and non-vesicular extracellular particles (NVEPs) were isolated from human plasma by differential centrifugation. Plasma was centrifuged at 1,000xg for 20 minutes to remove debris. Plasma was diluted 1:5 in PBS and was then centrifuged at 10,000xg for 40 minutes and the pellet (lEVs) was resuspended in resuspended in PBS containing 25 mM HEPES (pH 7.2) (PBS-H). The supernatant was then centrifuged at 167,000xg for 4 hours in a SW32 Ti swinging-bucket rotor (Beckman Coulter) and the pellet (sEVs) was resuspended in resuspended in PBS-H. Then the supernatant was ultracentrifuged at 167,000g for 16 hours. The resulting pellet (exomeres) was resuspended in PBS-H. The supernatant was ultracentrifuged at 367,000g for 16 hours and the resulting pellet was resuspended in PBS-H (supermeres).

### Transmission electron microscopy

Isolated extracellular vesicles and non-vesicular extracellular particles were imaged by transmission electron microscopy. Briefly, 10 μL of each sample was freshly thawed and adsorbed to glow-discharged carbon-coated 400 mesh copper grids by flotation for 2 min. Three consecutive drops of 1× Tris-buffered saline was prepared on Parafilm. Grids were washed by moving from one drop to another, with a flotation time of 10 seconds on each drop. The rinsed grids were then negatively stained with 1% uranyl acetate (UAT) with tylose (1% UAT in deionized water (dIH_2_O), double-filtered through a 0.22-μm filter). Grids were blotted, and then excess UAT was aspirated, leaving a thin layer of stain. Grids were imaged on a Hitachi 7600 TEM operating at 80 kV with an XR80 charge-coupled device (8 megapixels, AMT Imaging, Woburn, MA, USA).

### Atomic force microscopy

Isolated exomeres and supermeres were imaged by atomic force microscopy. Before applying the exomeres and supermeres samples, a freshly cleaved mica was treated with NiCl_2_ using an established protocol with some modifications^63^. Briefly, 20 μL of 100 mM NiCl_2_ were applied to a freshly cleaved mica disc, incubated for 1 minute, and rinsed with 200 μL of PBS for five times. The disc was then dried with filter paper. Then isolated exomeres and supermeres were diluted 100 times in PBS. Both samples were aliquoted in 5 μL each and stored at - 80°C before they use. The 5 μl aliquot was then diluted with 495 μL of buffer immediately before depositing onto the NiCl2 pretreated mica surface.

Experiments were performed in imaging buffer (1X PBS buffer) at room temperature (∼ 25 °C) using a commercial apparatus (MFP-3D Origin, Asylum Research, Oxford Instruments). Images were acquired in tapping mode using biolever mini cantilevers (AC40TS, Olympus) with nominal spring constant k ∼ 0.09 N/m. Images were taken in 512 × 512 resolution with a scan rate of 1 Hz. Care was taken to control the tip-sample force to be <100 pN. Prior to image analysis, images were flattened (1st order) to minimize background. Image analysis was done using the built-in particle analysis software (Asylum Research).

### Immunoblotting

Protein samples were measured by BCA assay (Catalog number 23227, Thermo Fisher). Equal amount of protein was loaded per well. The pellets were lysed in 10X RIPA buffer (Catalog number #9806, Cell Signaling) and protease inhibitors (Catalog number 78425, Thermo Fisher) for 15 min on ice. Samples were spun down for 15 min at 14,000 rpm in a tabletop centrifuge at 4°C. Fractions or lysates were mixed with 4× TGX sample buffer (Catalog number 1610747, Bio-Rad) under non-reducing conditions (for EV tetraspanin blotting) or under reducing conditions, boiled for 5 min at 95°C, and run into a PAGE gel electrophoresis on a 4%–20% Criterion TGX Stain-Free Precast gel (Catalog number 5678094, Bio-Rad). Proteins were transferred to a polyvinylidene fluoride (PVDF) membrane using the turbo transfer system (Bio-Rad) using the mix molecular weight program. After membranes were blocked for 1 h in 5% BSA in PBS + 0.05% Tween-20 (PBS-T), blots were incubated overnight at 4°C with the indicated primary antibodies in blocking buffer (Table 1). Then blots were washed 3 times with PBS-T and incubated for 1 h at room temperature with secondary antibodies (Table 1). After membranes were washed 3 times with PBS-T and developed with SuperSignal West Femto (Catalog number 34095, Thermo Fisher), SuperSignal West Atto Ultimate Sensitivity Substrate (Catalog number A38555, Thermo Fisher) or fluorescence on an iBright FL1500 (Thermo Fisher) in chemiluminescence or fluorescence mode.

### RNA isolation and qPCR from EVs and NVEPs

RNA isolation was performed using miRNeasy Mini Kit (Catalog number 217004, Qiagen) as per manufacturers protocol. Reverse transcription was performed using miRCURY LNA RT Kit (Catalog number 339340, Qiagen) according to the manufacturer’s instructions. RNA integrity was determined with Nanodrop (Nanodrop One, Thermo Fisher Scientific) and a 4150 TapeStation System (Agilent Technologies, Santa Clara, CA, USA). Quantitative real-time PCR (qRT-PCR) was performed using miRCURY LNA SYBR Green PCR Kit (Catalog number 339346, Qiagen) and the QuantStudio 6 Flex instrument (Applied Biosystems). Primers for hsa-miR-146a-5p (Catalog number 339306, GeneGlobe ID - YP00204688) and the C. elegans miR-39 spike-in control (Catalog number 339306, GeneGlobe ID YP00203952) were purchased from Qiagen. Relative miRNA levels from EVs and NVEPs were established against C. elegans miR-39 mimic spike-in control, using the ΔΔCq method.

### RNA extraction and qPCR from cells and tissues

Total RNA was extracted using RNeasy Mini kit according to the manufacturer’s protocol (Catalog number 74104, Qiagen). For cells, 1×10^6^ cells were lysed by adding 500 ml QIAzol lysis reagent. For tissues, 25 mg was used for RNA isolation. cDNA was synthesized with 0.5-1 μg of total RNA using High-Capacity cDNA Reverse Transcription Kit (Catalog number 4368813, Thermo Fisher) according to the manufacturer’s instructions. Quantitative real-time PCR (qRT-PCR) was performed using Ssoadvanced Universal SYBR (Catalog number 172527, Bio-Rad) following manufacturer’s instructions and the QuantStudio 6 Flex instrument (Applied Biosystems). All samples were assessed to the levels of Rsp18 expression as an internal control. qPCR data were assessed and reported according to the 2-ΔΔCt method.

### Nextflex Small RNA sequencing Library Generation

Small RNA sequencing libraries were generated from 22.5ng of Total RNA in 5ul of water using Nextflex Small RNA Sequencing kit V4 (revvity). No blockers were used, and all remaining steps of the protocol were followed as per manufacturer. Adapters were diluted by one-half and final indexed libraries were created by PCR from bead-cleaned cDNA using 20 cycles of amplification. Final libraries were characterized using high-sensitivity DNA Bioanalyzer chips. 1 ul of each library was used to create a pool and then sequenced on the ISEQ to determine mapping and to generate a normalized pool for sequencing on the Novaseq X plus sequencer (Illumina). The final equimolar pool was quantified by qPCR and High Sensitivity DNA D1000 on the Tapestation 4200 (Agilent). Sequencing Run was demultiplexed so that 50 read cycles and 8 indexed cycles for the i7 and i5 were generated for each fastq.

### NanoFlow Cytometry

Isolated EVs or NVEPs were diluted in 0.01 M HEPES buffer to a particle concentration of approximately 1× 10^10^ particles/ml for staining. Briefly 9 µL of the sample was incubated with 1 µL of the corresponding antibodies previously diluted in the dark for 30 minutes (Table 1). Then sample was diluted 100-fold prior to analysis in a nanoflow cytometer (Flow Nanoanalyzer, NanoFCM). Before analysis, the instrument was calibrated for particle concentration and for size distribution NanoFCM™ Silica Nanospheres Cocktail #1 with different sizes 68 to 155 nm (Catalog number S16M-Exo, NanoFCM). Samples were run and data collected for 1 minute with a pressure of 1.0 kPa. Data was analyzed using data were analyzed by the NanoFCM Software V1.17 (NanoFCM).

### Microfluidic resistive pulse sensing

Microfluidics resistive pulse sensing measurements were performed with the nCS1 instrument (Spectradyne, Torrance, CA). Isolated extracellular vesicle samples were diluted 1:100 in 1% Tween 20 in 1× PBS (PBST) and loaded onto polydimethylsiloxane cartridges (diameter range 65 nm to 400 nm). A different cartridge was used for each sample and replicate. Approximately 10 µL of the diluted sample (1:100) was used, and events were recorded for each sample. Results were analyzed using the nCS1 Data Analyzer (Spectradyne, Torrance, CA).

### Nanoparticle Tracking Analysis

Exosome-containing fractions were diluted into 10-nm filtered PBS and examined for concentration of exosome-sized vesicles and particles by NTA using a Particle Metrix Particle Metrix ZetaView Evolution, according to the manufacturer’s instructions.

### Human Tetraspanin ELISA

The human tetraspanin ELISA (Catalog number Atlas Human EV ELISA Kit, Everest Biolabs), was run as per the manufacturer’s instructions. Briefly, EV standards and samples were diluted (1:50 for human plasma EVs or NVEPs, 1:6 for cell culture EVs or NVEPs) in sample diluent buffer and the antibody cocktail was added at 1:1 ratio, then incubated overnight at 4°C. Next plates were washed with 1x wash buffer five times and incubated with TMB substrate solution for 10 minutes in the dark on a plate shaker at 600 rpm. Then the stop solution was added to each well and subjected to shaking for 1 minute at 300 rpm. Absorbance was measured at 450 nm using a plate reader (SpectraMax® iD3, Molecular Devices). All samples were run in two technical replicates and three biological replicates.

### Human APOA ELISA

Atlas APOA ELISA Kit (Catalog number Atlas APOA ELISA Kit, Everest Biolabs) was run as per the manufacturer’s instructions. Briefly, standards and samples were diluted 1:60 in sample diluent buffer and the antibody cocktail was added at 1:1 ratio, then incubated overnight at 4°C. Next, plates were washed with 1x wash buffer five times and incubated with TMB substrate solution for 10 minutes in the dark on a plate shaker at 600 rpm. Then the stop solution was added to each well and placed on shaker for 1 minute at 300 rpm. Absorbance was measured at 450 nm using a plate reader (SpectraMax iD3, Molecular Devices). All samples were run in two technical replicates and three biological replicates.

### Immunofluorescence assays in cells

HMC3 cells or iTF-Microglia cells plated on chamber slides were washed with PBS and fixed with 4% cold paraformaldehyde (Catalog number sc-281692, Santacruz Biotechnology). The cells were permeabilized with 0.1% Triton X-100 (Catalog number X100-500ML, Sigma) for 10 minutes at room temperature (RT) and blocked with 5% bovine serum albumin (Catalog number A7906-500G, Sigma) for 1 hour at RT. Primary antibodies were incubated overnight at 4°C diluted in 5%BSA (Table 1). Cells were washed with 1xPBS (Catalog number 10010049, Thermo Scientific) three times (5 minutes each wash). The corresponding secondary antibodies were incubated for 1 hour at RT (Table 1) and DAPI (Catalog number 62248, Thermo Fisher) was used to stain the nuclei incubated for five minutes. Cells were washed with 1xPBS (Catalog number 10010049, Thermo Scientific) three times (5 minutes each wash) before the slides were covered with coverslips using Vectashield (Catalog number H-1700-10, Vector Laboratories). Images were acquire using a Leica SP8 microscope.

### SEVEN Assay

#### Lectin and antibody conjugation with Fluorophores and characterization of fluorescent probes

SNA lectin was conjugated with Alexa Fluor 647 N-hydroxysuccinimidyl (NHS) ester dye (AF647; Invitrogen, Cat# A20006) while αCD9, αCD63, and αCD81 antibodies were conjugated with either AF647 or Biotium CF568 succinimidyl ester dye (CF568; Fisher Scientific, Cat# NC154276) according to a previously established protocol^64^. The degree of labeling was quantified using a NanoDrop 1000 spectrophotometer (Thermo Fisher Scientific), with a typical degree of labeling ranging from 1.0 to 2.0. The maximum dark time and average number of localizations per molecule for each fluorescent reporter were evaluated as previously described^30,65^. The average number of localizations per fluorescent probe (α) was: 6 for SNA-AF647; 10 for anti-human αCD9, αCD63, and αCD81antibodies-AF647 and anti-mouse αCD9, αCD63, and αCD81 with anti-human αCD81 antibodies-AF647; 11 for anti-human αCD9, αCD63, and αCD81 antibodies-CF658; and 11 for anti-ApoE antibody-AF647.

#### Analytical protocol for SEVEN

We followed a previously established and validated analytical protocol^30^. Briefly, 25 mm glass coverslips (#1.5H, Catalog number NC9560650, Thermo Fisher Scientific) were cleaned and functionalized with MCP4 polymer (Catalog number MCP4-KIT, Lucidant Polymers). A 0.5 μL volume of one of the following reagents was applied to the center of each coverslip for target capture: (1) SNA, (2) a mixture of anti-human TSPAN antibodies (αCD9, αCD63, and αCD81), (3) anti-human αCD81 antibody, and (4) anti-human αApoE antibody, all prepared in PBS supplemented with 1% glycerol (Table 1). Final concentrations were 2.5 mg/mL for SNA, 0.17 mg/mL for each of the anti-human TSPAN antibodies in the mixture and 0.5 mg/mL for anti-human αCD81 and ApoE antibodies. After 4 hours of incubation at room temperature in a humidity-controlled environment, the surfaces were washed, blocked, and incubated with EV- or NVEP- containing samples. Due to differences in EV/NVEP concentrations across sample types, different dilutions were applied.

For SNA enrichment, we used reconstitution buffer: 0.025% (v/v) Tween-20 in phosphate-buffered saline (PBS) (0.025% (v/v) PBS-T) supplemented with 10 mM HEPES and 0.1 mM Ca²⁺. For SNA-enriched NVEP detection from isolated microglia fractions, samples were diluted 1,000-fold; for SNA-enriched NVEP detection from crude plasma and isolated plasma fractions, samples were diluted 10,000-fold; and for SNA-enriched EV detection from crude plasma and isolated plasma fractions, samples were diluted 1:100 fold. Samples were incubated overnight at room temperature with gentle shaking and subsequently washed. Samples were pre-fixed in 4% (w/v) paraformaldehyde (Catalog number 157-8, Electron Microscopy Sciences) for 30 minutes at room temperature. After quenching with 25 mM glycine in PBS for 10 minutes and washing with PBS, fluorescent staining was performed using (1) 10 nM SNA-AF647 for detection of NVEPs or (2) or a mixture of 10 nM αCD9-AF647, 10 nM αCD63-AF647, and 10 nM αCD81-AF647 for detection of EVs in 2% (w/v) BSA in 0.025% (v/v) PBS-T. Following staining, samples were washed, post-fixed in 4% paraformaldehyde in PBS for 15 minutes at room temperature, quenched with 25 mM glycine in PBS for 10 minutes, and washed again.

For TSPAN and ApoE enrichment, we used reconstitution buffer: 0.025% (v/v) PBS-T. For TSPAN-enriched EV detection from isolated microglia fractions, samples were diluted 10,000-fold; for TSPAN-enriched EV detection from isolated plasma fractions, samples were diluted 1,000-fold; for TSPAN-enriched EV detection from crude plasma, samples were diluted 5,000-fold; for ApoE-enriched EV and LPP detection from crude plasma, samples were diluted 500-fold. As before, samples were incubated overnight at room temperature with gentle shaking and subsequently washed. Samples captured with αTSPAN antibodies were stained with a mixture of αCD9-AF647, αCD63-AF647, and αCD81-AF647, each at 10 nM (for EV detection). Samples captured with ApoE antibody were stained with 10 nM ApoE antibody labeled with AF647 (for LPP detection) and a mixture of 10 nM αCD9-CF568, αCD63-CF568, and αCD81-CF568 antibodies (for EV detection). Staining was performed in 2% (w/v) BSA in 0.025% (v/v) PBS-T. After staining, samples were washed, fixed in 4% (w/v) paraformaldehyde, 0.2% (w/v) glutaraldehyde in PBS for 30 minutes at room temperature, quenched with 25 mM glycine in PBS for 10 minutes, and washed again.

For hCD81 enrichment, we used reconstitution buffer: 0.025% (v/v) PBS-T. sEVs isolated from mice plasma and brain tissue were directly applied onto the functionalized coverslips without further dilution. After overnight incubation at room temperature (with gentle shaking) and wash, samples were stained with a mixture of mouse αCD9-AF647, αCD63-AF647, αCD81-AF647, and human αCD81-AF647, each at 10nM. Staining was performed in 2% (w/v) BSA in 0.025% (v/v) PBS-T. After staining, samples were washed, fixed in 4% (w/v) paraformaldehyde, 0.2% (w/v) glutaraldehyde in PBS for 30 minutes at room temperature, quenched with 25 mM glycine in PBS for 10 minutes, and washed again.

### Imaging and Analysis

For imaging, samples were placed in Attofluor cell chambers (Thermo Fisher Scientific, Cat# A7816) and imaged using a 3D N-STORM super-resolution microscope (Nikon Instruments, Melville, NY, USA) in dSTORM buffer^66^. A 640 nm laser with an excitation power of 124 mW, and 561 nm laser with an excitation power of 110 mW, where noted, were employed for dSTORM imaging. A total of 25,000 frames were captured at a 10 ms exposure time for each region of interest (ROI), measuring 41 × 41 μm (256 × 256 pixels). For two-color imaging, 561 nm laser acquisition was performed following 640 nm laser acquisition. Image acquisition was done using NIS-Elements software (Nikon Instruments, version 5.21.01 and 5.41.0).

After peak localization with NIS-Elements, analysis of the raw SMLM images was performed using MATLAB R2024b using Voronoi tessellation-based clustering^30^. Two-color images were aligned as reported before^30^. The following parameters were used for analysis of human plasma and isolated microglia fractions: for small clusters (20-100 nm in diameter), the minimum localizations per cluster were 2xα, while the maximum localizations per cluster were 2,000. For large clusters (above 100 nm), the minimum localizations per cluster were 100, while the maximum localizations per cluster were 15,000. To avoid clustering artifacts due to high spinning speeds, for SNA enrichment/SNA detection data, we limited the size to 250 nm. For mouse plasma and tissue imaging, the minimum localizations per small cluster (30-100 nm) were set at 2.5xα to avoid any potential fragments, while large clusters had the same settings.

eSRRF image analysis was conducted using a custom macro on ImageJ (version 1.535; National Institutes of Health, USA) with previously optimized parameters^67^. The eSRRF processed images were binarized and thresholded using an optimized custom absolute threshold: 0.3 for tissue EVs and 0.4 for plasma EVs (values were optimized to yield minimal false positive signal and maximal overlap between channels). Subsequently, images were aligned with the corresponding 640 nm SMLM channel. SMLM data (red channel) for EVs that were positive for mNeonGreen (mNG, green channel) were reported.

#### Isolation of primary mouse microglia

Whole brains were collected on an ice-cold plastic dish from mice after euthanasia (2-3 months old mice, mixed gender). Tissues were mechanically homogenized and digested with 450 U/mL of collagenase I (10000 U/ml) (Catalog number C0130, Sigma), 125 U/mL of collagenase XI (12500 U/ml) (Catalog number C7657, Sigma), 60 U/mL of DNase I (Catalog number D5319, Sigma) and 60 U/mL of hyaluronidase (12000 U/ml) (Catalog number H3506, Sigma) for 15 minutes at 37C°. Following digestion, tissue suspensions were filtered through a 70-µm cell strainer, washed, and centrifuged at 300×g for 10 min to obtain single cells. The resulting homogenate was centrifuged again at 600×g for 6 min at room temperature (RT). The supernatant was carefully removed, and the pellet was resuspended in 6 mL of sterile 70% standard isotonic percoll (SIP). For density gradient separation, the resuspended homogenate was transferred to a sterile 15 mL polypropylene conical tube. Layers were prepared by carefully overlaying 3 mL of sterile 50% SIP on top of the 70% layer, followed by 3 mL of sterile 35% SIP, and finally 2 mL of sterile 1× DPBS. Prepared gradients were centrifuged at 2000 × g for 20 min at RT with no brake. After centrifugation, three distinct layers formed. The top myelin layer was discarded, and cells were collected from the 50%–35% interfaces into a separate tube. Collected cells were washed in sterile 1× DPBS and centrifuged at 600 × g for 6 min at RT to remove residual Percoll. Isolated microglia were immediately processed for downstream applications.

#### Flow cytometry

Isolated primary microglia were stained with anti-CD45-BV711 (clone 30-F11, BioLegend), anti-CD11b-BV785 (clone M1/70, BioLegend), and Live/Dead Fixable Aqua viability dye (Thermo Fisher). Cells were first incubated with the Live/Dead dye in PBS for 20 min on ice, washed twice with PBS, and subsequently stained with surface antibodies for 30 min on ice in the dark. After staining, cells were washed twice with PBS and resuspended in FACS buffer (PBS supplemented with 2% FBS and 2 mM EDTA) for acquisition. Data were acquired on a BD FACS Aria III flow cytometer (BD Biosciences) and analyzed using FlowJo software version 11. Microglia were identified as CD11b⁺CD45low events after exclusion of doublets and dead cells. Representative gating controls included fluorescence and unstained controls. The gating strategy can be found in Extended data Fig. 6b.

#### Immunocapture of microglia-derived extracellular vesicles from mouse plasma

Plasma was collected in EDTA tubes (Catalog number 366643, BD) and centrifuged at 2000xg for 10 min at 4°C. EVs were isolated from plasma by size exclusion chromatography (Catalog number qEV35, Izon). Plasma-derived extracellular vesicles were incubated with 80 µl of pre-washed anti-streptavidin magnetic beads (Catalog number 88817, Thermo Fisher Scientific) in 0.01% PBS-T on a gentle rotator shaker overnight at 4°C. Immunoprecipitates were washed three times with 1 ml 0.01 %PBS-T on a DynaMag Spin Magnet. For protein extraction from EVs the beads with bound EVs were resuspended in 25 µl of lysis buffer for Immunoblotting or 500 µl of trizol for RNA isolation. The mixture was incubated for 10 min at 4°C, after which the beads were removed using the magnet.

#### RNA-Sequencing Data Analysis

The raw sequence image files from the Illumina NovaSeq 6000 (bcl files) were converted to the fastq format using bcltofastq v. 2.19.1.403 and checked for quality to ensure the quality scores did not deteriorate at the read ends. All samples were processed using the exceRpt small RNA pipeline (PMID: 30956140). Briefly, reads in fastq files were trimmed to remove TruSeq adapters from the 3′ end using cutadapt. Reads shorter than 15 nucleotides after adapter trimming were discarded. After trimming, the reads were aligned to the human genome (GRCh38) using STAR. Since multi-mappers are common with small RNA (short read lengths), biotype was assigned from the STAR aligned BAM file in the priority order used in the exceRpt pipeline: miRNA, YRNA, tRNA, piRNA, and lastly protein-coding. Raw gene counts tables were then loaded into R (v. 4.5.1) for further analysis.

#### Differential Expression Analysis

Differential expression was conducted using the DESeq2 package (version 1.38.3) in R for all sequences that had expression levels > 10 read counts in at least 50% of samples, in each comparison. The raw read counts for the samples were normalized using the median ratio method (default in DESeq2). The Benjamini–Hochberg method was used for reported adjusted p-values.

#### Data preparation and statistical Analysis

Statistical analysis was performed using GraphPad Prism 10 software (version 10.3.1, GraphPad) and MATLAB. Two-tailed student’s t-test were used for statistical comparison between two groups, with *p* values of 0.05 or less considered significant (\**p*<0.05, \*\**p*<0.01, \*\*\**p*<0.001, \*\*\*\**p*<0.0001). If other tests were used, they were indicated in the figure legend. Figure diagrams and schematics were generated using BioRender with a permissive license. All figures were finalized using Adobe Illustrator 2022 (version 26.5.3; Adobe, San Jose, CA, USA).

### Data Availability Statement

Data is available in the main text and the supplemental information.

### Code Availability

The source code used for analysis of small RNA sequencing is available at https://github.com/gersteinlab/exceRpt.

The source code used in this study for SEVEN analysis is available at https://github.com/andronovl/SharpViSu ^68^ with parameters indicated^67^.

## Acknowledgments

The authors would like to thank all members of the Das, Van Keuren-Jensen, and Talisman laboratories. We also thank David Palmlund and Particle metrix for the technical support. We also thank the Johns Hopkins Microscope Facility for the technical support and I. Talisman for manuscript editing. The authors gratefully acknowledge support from the flow cytometry core at the Massachusetts General Hospital - Simches Research Center. This work was supported by the National Institutes of Health grants UG3/UH3 TR002878 (KVKJ, TJT, SD) and R01 AA028549 (TJT), Diabetes Prevention Risk Omics Metabolism and Therapy of Diabetes (PROMT) Interdisciplinary Training Program T32 DK131943 (BP).

## Authors contributions

M.G.C. performed and analyzed experiments. C.L. B.P., N.J. performed SEVEN experiments. C.L. B.P., N.J. A.S., C.T. analyzed SEVEN experiments. B.K, O.D.W., S.H., J.F., and B.S performed and analyzed selected experiments. B.M. prepared the libraries for sequencing. E.A. performed the bioinformatic analyses. J.F. assisted with Nanoflow cytometry experiments. M.G.C., C.L., S.D., K.VK. J and T.J.T contributed to the experimental design. KP.S. supervised AFM experiments. K.VK. J supervised the bioinformatic analysis. S.D., K.VK. J, and T.J.T. directed the research. M.G.C. wrote the paper with assistance from all other authors.

